# Mapping *in vivo* genetic interactomics through Cpf1 crRNA array screening

**DOI:** 10.1101/153486

**Authors:** Ryan D. Chow, Guangchuan Wang, Adan Codina, Lupeng Ye, Sidi Chen

## Abstract

Genetic interactions lay the foundation of biological networks in virtually all organisms. Due to the complexity of mammalian genomes and cellular architectures, unbiased mapping of genetic interactions *in vivo* is challenging. Cpf1 is a single effector RNA-guided nuclease that enables multiplexed genome editing using crRNA arrays. Here we designed a Cpf1 crRNA array library targeting all pairwise permutations of the most significantly mutated non-oncogenes, and performed double knockout screens in mice using a model of malignant transformation as well as a model of metastasis. CrRNA array sequencing revealed a quantitative landscape of all single and double knockouts. Enrichment, synergy and clonal analyses identified many unpredicted drivers and co-drivers of transformation and metastasis, with epigenetic factors as hubs of these highly connected networks. Our study demonstrates a powerful yet simple approach for *in vivo* mapping of unbiased genetic interactomes in mammalian species at a phenotypic level.

## Introduction

Genetic interactions lay the foundation of virtually all biological systems ^1,2^. With rare exceptions, every gene interacts with one or more other genes, forming highly complex and dynamic networks ^3, 4^. The nature of genetic interactions include physical interactions, functional redundancy, enhancer, suppressor, or synthetic lethality ^4^. Such interactions are the cornerstones of biological processes such as embryonic development, homeostatic regulation, immune responses, nervous system function and behavior, and evolution ^5^. Perturbation or misregulation of genetic interactions in the germ line can lead to failures in development, physiological malfunction, autoimmunity, neurological disorders, and many forms of genetic diseases ^6,7^. Disruption of the genetic networks in somatic cells can lead to malignant cellular behaviors such as uncontrolled growth, driving the development of cancer ^8,9^.

The study of genetic interactions evolved over a century, originating in the era of classical genetics ^10^. In essence, how two genes interact can be studied by examining the phenotypes of double mutants as compared to single mutants ^11^. This concept of epistasis has guided the conceptualization and subsequent discovery of countless important pathways, and has become the gold standard for determining downstream and upstream regulation in genetic analysis ^2, 12^. For instance, synthetic lethality has been investigated in animal development and cancer therapeutics ^8,13^. Classical approaches such as genome-wide association studies (GWAS) and quantitative trait loci (QTL) mapping have been extensively employed to study complex phenotypes that involve multiple genes ^12,14^.

The discovery and characterization of the type V CRISPR system, Cpf1 (CRISPR from *Prevotella* and *Francisella*) has enabled rapid genome editing of multiple loci in the same cell ^15–19^. Cpf1 is a single component RNA-guided nuclease that can mediate target cleavage with a single crRNA ^17,19^. Compared to Cas9, Cpf1 does not require a tracrRNA, which greatly simplifies multiplexed genome editing of two or more loci simultaneously by using a string of crRNAs targeting different genes ^17,19^. These characteristics led us to hypothesize that Cpf1 might be an ideal system for high-throughput higher dimensional screens in mammalian species, with substantial advantages in library design and readout when compared to Cas9-based approaches. We designed a Cpf1 crRNA array library targeting a set of the most significantly mutated cancer genes and performed an unbiased screen on two different mouse models, one studying early-stage tumorigenesis and the second studying cancer metastasis, identifying many unpredicted gene pairs. Thus, Cpf1 screening is a powerful approach to systematically quantify genetic interactions and identify new synergistic combinations. Unlike with Cas9-based strategies, due to the simple expansion of crRNA arrays, this approach can be readily extended to perform triple-, quadruple- or higher dimensional screens *in vivo*.

## Results

### Enabling one-step double knockout screening with a Cpf1 crRNA array library

To establish a lentiviral system for CRISPR/Cpf1-mediated genetic screening, we first generated a human-codon-optimized LbCpf1 expression vector (pLenti-EFS-Cpf1-blast, LentiCpf1 for short) and a crRNA expression vector (pLenti-U6-DR-crRNA-puro, Lenti-U6-crRNA for short) (Fig. 1a). In order to facilitate direct and targeted double knockout studies using a single crRNA array, we designed oligos with a 5’ homology arm to the base vector, followed by a crRNA, the direct repeat (DR) sequence for Cpf1, a second crRNA, a U6 terminator, and finally a 3’ homology arm (cr1-DR-cr2). As the oligos each contain two crRNAs, we termed these constructs crRNA arrays. Linearization of the Lenti-U6-crRNA vector enables one-step cloning of the crRNA array into the vector by Gibson assembly, producing the double knockout crRNA array expression vector (pLenti-U6-DR-cr1-DR-cr2-puro) (Fig. 1b). We first tested these constructs for their ability to induce double knockouts in a murine cancer cell line (KPD) ^20,21^ *in vitro*. After infection with LentiCpf1, we then transduced the cells with lentiviruses carrying a crRNA array targeting *Pten* and *Nf1* (Fig. S1a). To confirm whether Cpf1 can mediate mutagenesis regardless of the position of each crRNA within the array ^19^, we generated two permutations of the *Pten* and *Nf1* crRNA array (crPten.crNf1 and crNf1.crPten, all with 20nt spacers) (Fig. S1a), and found that both crPten.crNf1 and crNf1.crPten crRNA arrays generated indels at both loci in Cpf1+ KPD cells (Fig. S1b). These data confirmed that a single crRNA array can be used in conjunction with CRISPR-Cpf1 to mediate simultaneous knockout of two genes in mammalian cells.

**Figure 1:**
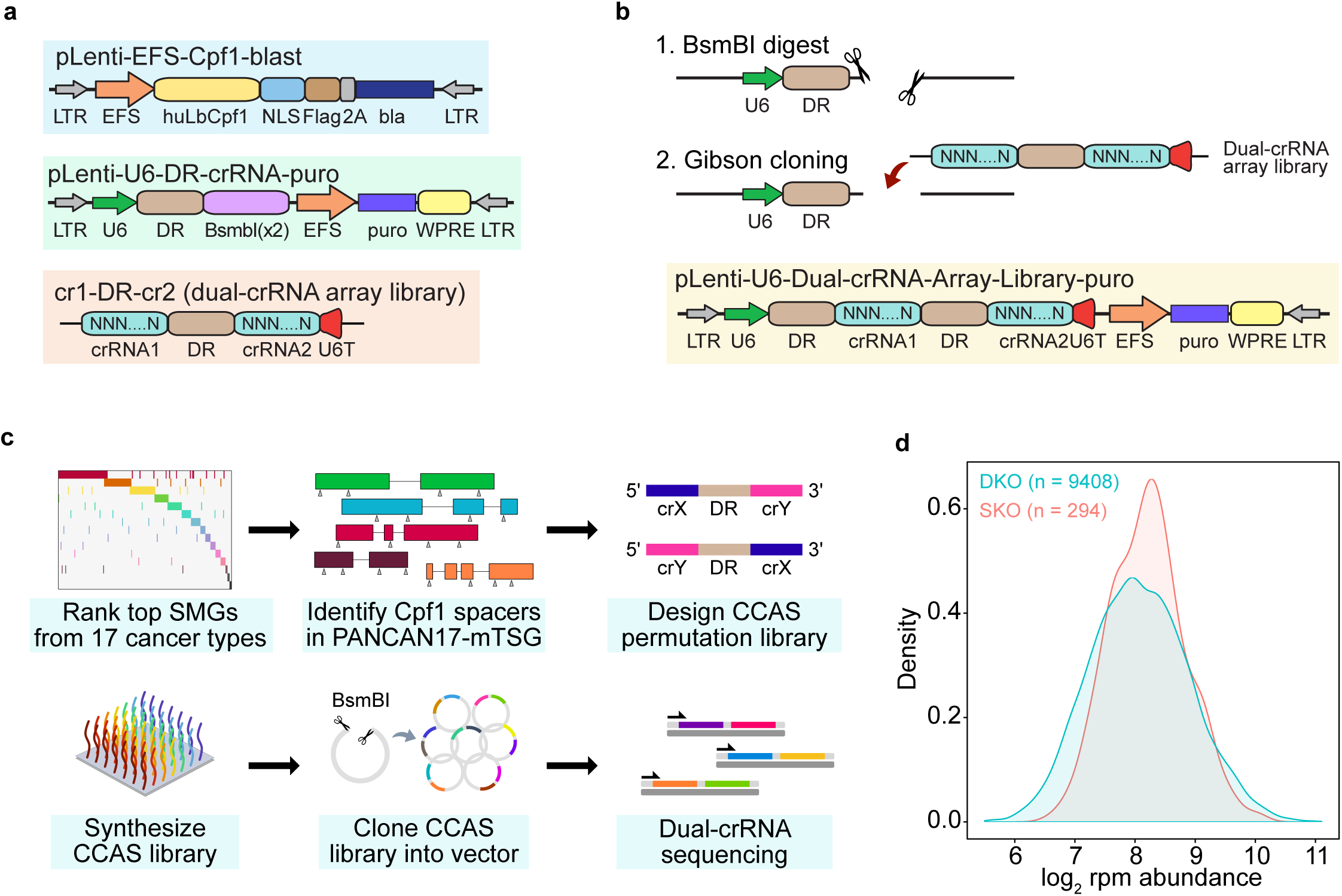
Enabling one-step double knockout screening with a Cpf1 crRNA array library. **a.** Schematic maps of the constructs for one-step double knockout screens by CRISPR-Cpf1. We generated a pLenti-EFS-Cpf1-blast vector (blue box), which constitutively expresses a humanized form of *Lachnospiraceae bacterium* Cpf1 (LbCpf1); transduced cells can be selected by blasticidin. We also generated a pLenti-U6-DRcrRNA-puro vector (green box), which contains the direct repeat (DR) sequence of Cpf1 and two BsmBI restriction sites for one-step cloning of crRNA arrays; puromycin treatment enables the selection of cells that have been transduced. The structure of the crRNA array library for cloning into the base vector is also shown (orange box). Each crRNA array is comprised of a 5’ homology arm to the base vector, followed by the first crRNA, the direct repeat (DR) sequence for Cpf1, the second crRNA, a U6 terminator sequence, and a 3’ homology arm. **b**. Schematic of the cloning strategy for double knockout screens by CRISPR-Cpf1. Incorporating a crRNA array library into the base vector simply requires BsmBI linearization followed by Gibson assembly, thereby producing a lentiviral version of the library (pLenti-U6-DR-cr1(N20)-DR-cr2(N20)-puro, yellow box). This one-step cloning procedure greatly simplifies library construction for high-dimension genetic screens. **c**. Schematic describing the design and synthesis of the Cpf1 double knockout (CCAS) library for identifying synergistic drivers of tumorigenesis. We first identified the top 50 tumor suppressors (TSGs) based on an unbiased pan-cancer analysis of 17 cancer types from the TCGA (PANCAN17-TSG50). 49 of these 50 TSGs had a corresponding mouse orthologs (PANCAN17-mTSG). All possible Cpf1 spacer sequences within these genes were identified and 2 were chosen for each gene. The selection of crRNAs was based on two scoring criteria: 1) high genome-wide mapping specificity and 2) a low number of consecutive thymidines, since long stretches of thymidines will terminate U6 transcription. With these 98 crRNAs and 3 additional non-targeting control (NTC) crRNAs, a library was designed containing 9,705 permutations of two crRNAs each (CCAS library). After pooled oligo synthesis, the PANCAN17-mTSG CCAS library was cloned into the base vector, and the plasmid crRNA array representation was subsequently readout by deep-sequencing the crRNA expression cassette. **d**. Density plots showing the distribution of CCAS crRNA array abundance in terms of log2 reads per million (rpm). Of the 9,705 total crRNA arrays in the library, 9,408 were comprised of two gene-targeting crRNAs (double knockout, or DKO; teal), while 294 contained one gene-targeting crRNA and one NTC crRNA (single knockout, or SKO; orange). The remaining 3 crRNA arrays were controls, with two NTC crRNAs in the crRNA array (NTCNTC, not shown). The library-wide abundance of both DKO and SKO crRNA arrays followed a log-normal distribution, demonstrating relatively even coverage of the CCAS plasmid library.

To investigate whether Cpf1 multiplex gene targeting could be utilized for multidimensional genetic interaction screens, we next sought to develop a library for Cpf1 crRNA array screening (CCAS library). Considering the resolution of library complexity under *in vivo* cellular dynamics, we aimed to design a focused CCAS library of the top 50 significantly mutated genes (SMGs) that are not oncogenes, with the vast majority being established or putative tumor suppressor genes (TSGs) identified through analysis of 17 different cancer types from The Cancer Genome Atlas (TCGA) (Methods). We termed the resultant gene set as PANCAN17- TSG50. (Fig. 1c; Table S1) ^22–26^. 49 of the PANCAN17-TSG50 genes had corresponding mouse orthologs (PANCAN17-mTSG), and were thus included in the CCAS library. We identified all possible Cpf1 spacer sequences within PANCAN17-mTSG and subsequently chose 2 crRNAs for each gene. The selection of crRNAs was based on two scoring criteria: 1) high genome-wide mapping specificity and 2) a low number of consecutive thymidines, since long stretches of thymidines will terminate U6 transcription (crRNA scoring data not shown due to size, available upon request).

Compiling these 98 gene-targeting crRNAs and 3 additional non-targeting control (NTC) crRNAs, we designed a crRNA array library containing 9,705 permutations of two crRNAs each (Table S2). Of the 9,705 total crRNA arrays in the library, 9,408 were comprised of two gene-targeting crRNAs (double knockout, or DKO), while 294 contained one gene-targeting crRNA and one NTC crRNA (single knockout, or SKO). The remaining 3 crRNA arrays were dedicated controls, with two different NTC crRNAs in the crRNA array (NTC-NTC). After pooled oligo synthesis, we cloned the PANCAN17-mTSG CCAS library into the base vector, and subsequently readout the plasmid crRNA array representation by deep-sequencing the crRNA expression cassette. All 9,705/9,705 (100%) of the designed crRNA arrays were successfully cloned (Figures 1d, S2b; Tables S3-4). Analysis of each crRNA array within the CCAS library revealed that the relative abundances of both DKO and SKO crRNA arrays approximated a log-normal distribution, demonstrating even coverage of the CCAS library revealed that the relative abundances of both DKO and SKO crRNA arays approximated log-normal distribution, demonstrating even coverage of the CCAS library (Fig. 1d). We generated lentiviral pools from the CCAS plasmid library for subsequent high-throughput double-mutagenesis and genetic interaction screens.

### Library-scale Cpf1 crRNA array screen in a mouse model of early tumorigenesis

To perform an *in vivo* Cpf1 screen, we first utilized a mouse model of malignant transformation and early stage tumorigenesis. We transduced an immortalized murine cell line with low tumorigenicity (clone IM) ^27^ with LentiCpf1 and then with the CCAS lentiviral pool. The library transduction was performed with four infection replicates at high coverage (~2,000x coverage for each replicate) and low multiplicity of infection (MOI, ≤ 0.2) to ensure the vast majority of cells would only carry one provirus integrant (Methods) (Fig. 2a). After 7 days of puromycin selection, only CCAS-virus infected cells survived, comprising a mixture of various double mutants (termed CCAS-treated cells hereafter). In parallel, we infected another group of Cpf1+ cells with lentiviruses carrying the empty vector. We then subcutaneously injected the virus-treated cell populations into nude mice (CCAS, n = 10 mice; vector, n = 4 mice). By 45 days post-injection (dpi), CCAS-treated cells had given rise to significantly larger tumors than vector-treated cells (*p* = 0.0223, by two-sided t-test) (Fig. 2b). This trend continued through the duration of the experiment (46.5 dpi, *p* = 0.0017). A select fraction of tumors derived from CCAS-treated cells were harvested and sectioned for histological analysis, together with the small nodules derived from vector-treated cells (Fig. 2c).

**Figure 2:**
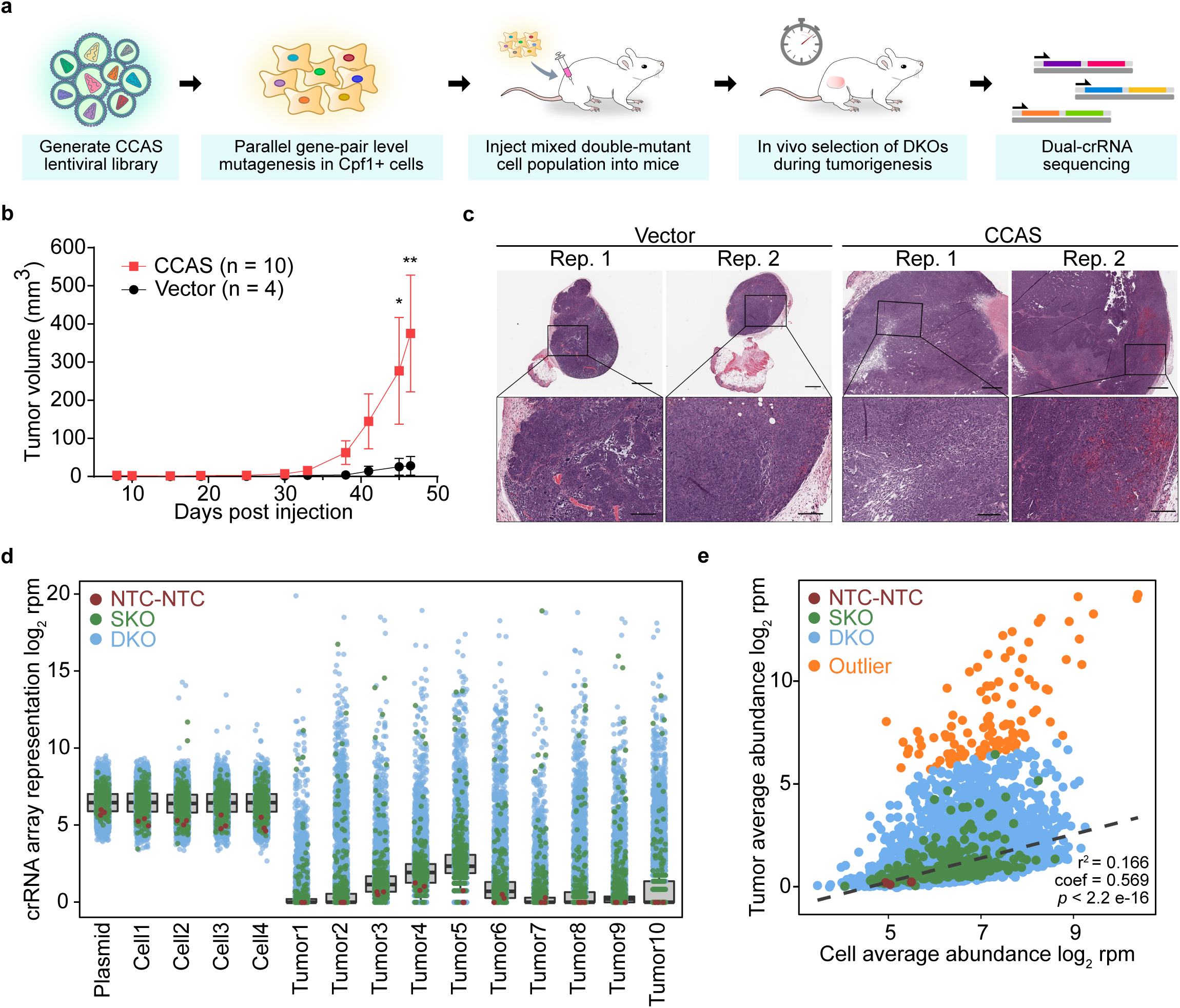
Library-scale Cpf1 crRNA array screen in a mouse model of early tumorigenesis. **a**. Schematic of the experimental approach for Cpf1-mediated double knockout screens to identify synergistic drivers of tumorigenesis in a transplant model. We generated lentiviral pools from the CCAS plasmid library, and subsequently infected Cpf1+ IM cells to perform massively parallel gene-pair level mutagenesis. The mixed double mutant cell population (CCAS-treated cells), or vector-treated control cells were then injected subcutaneously into nude mice (n = 10 and n = 4, respectively). After 6.5 weeks, genomic DNA was extracted from the injection site and subjected to crRNA array sequencing. **b.** Tumor growth curves of CCAS-treated (red, n = 10) and vector-treated cells (black, n = 4) *in vivo*. As expected, vector treated cells were lowly tumorigenic, and the population of mixed double mutants (CCAS-treated cells) were highly tumorigenic. By 45 days post injection (dpi), tumors derived from CCAS-treated cells were significantly larger than those by vector-treated cells (* *p* < 0.05, ** *p* < 0.01, two-sided t-test). **c**. Histological sections of tumors derived from vector-treated and CCAS-treated cells, stained by hematoxylin and eosin. Two representative tumors are shown from each group. Images in each row were taken at the same magnification (top row, scale bar = 500 μm; bottom row, scale bar = 200 μm). CCAS-treated cells gave rise to much larger tumors than vector-treated cells. **d.** Dot-boxplot depicting the overall representation of the CCAS library, in terms of log_2_ rpm abundance. The plasmid library, 4 pre-injection cell pools, and 10 tumor samples were sequenced. NTC-NTC controls are shown in red, SKO crRNA arrays in green, and DKO crRNA arrays in blue. Whereas plasmid and cell samples exhibited lognormal representation of the CCAS library, tumor samples showed strong enrichment of specific SKO and DKO crRNA arrays. Notably, NTC-NTC crRNA arrays were consistently found at low abundance in all tumor samples. **e.** Scatterplot comparing average log_2_ rpm abundance of all CCAS crRNA arrays in cells and in tumors. As above, NTC-NTC controls are shown in red, SKO crRNA arrays in green, and DKO crRNA arrays in blue. The linear regression line is shown, demonstrating the log-linear relationship of most crRNA arrays between tumors and cells (r^2^ = 0.166, coefficient = 0.569, *p* < 2.2 e-16 by F-test). There were numerous outliers (orange) (Bonferroni adjusted *p* < 0.05), indicating that specific crRNA arrays had undergone positive selection *in vivo*. See Fig. S3 for the individual tumor comparisons, with outliers labeled.

To unveil the genetic interactions that had driven rapid tumor growth upon Cpf1-mediated mutagenesis, we performed crRNA array sequencing on genomic DNA from CCAS tumors (n = 10) and pre-injection cell pools (n = 4) (Methods). Whereas plasmid and cell samples were highly correlated with one another, tumor samples were more correlated with other tumors (Fig. S2a). All plasmid and cell samples contained 100% of CCAS crRNA arrays, while tumor samples exhibited significantly lower crRNA array library diversity (mean ± SEM = 37.0% ±10.5%; *p* = 2.02 e-4 compared to plasmid and cells, t-test) (Fig. S2b, Table S5). Furthermore, while plasmid and cell samples exhibited robust lognormal representation of the CCAS library (Fig. S2c), tumor samples showed strong enrichment of specific SKO and DKO crRNA arrays (Fig. S2c, Fig. 2d; Table S4). Of note, the 3 NTC-NTC controls in the CCAS library were consistently found at abundances similar to one another within each of the plasmid and cell samples. All NTC-NTCs were found at low abundance across all tumor samples (average log2 rpm abundance = 0.224 ± 0.108), suggesting non-mutagenized cells do not have a selective advantage in tumorigenesis and that additional genetic perturbation is needed to drive rapid tumor growth *in vivo*. As a global comparison of all crRNA arrays, while the mean abundance in tumor samples correlated with the mean abundance in cell samples in a log-linear manner (regression r^2^ = 0.166, coefficient = 0.569, *p* < 2.2 e-16 by F-test), a population of crRNA arrays were outliers (Outlier test, Bonferroni adjusted *p* < 0.05), indicating that specific crRNA arrays had undergone positive selection *in vivo* (Fig. 2e, Table S6). This trend was consistent across the individual tumors (*p* < 2.2 e-16 by F-test for all individual tumors, average number of outliers compared to cells = 102.1 ± 3.671 crRNA arrays) (Fig. S3a-b). Taken together, these data suggest that a subgroup of crRNA arrays was enriched in tumors, indicating that a select number of mutant clones had significantly expanded *in vivo*.

### Enrichment analysis of single knockout and double knockout crRNA arrays

To further investigate the specific genetic interactions that had driven early stage tumorigenesis in CCAS-treated cells, we next examined the distribution of raw crRNA array abundance within each sample (Methods). Within each tumor, we observed specific crRNA arrays that were heavily enriched by several orders of magnitude, suggesting that these mutant clones had undergone potent positive selection (Figures 3a, S2d). For example, in Tumor 1, crCasp8.crApc was by far the most abundant crRNA array, dwarfing all other crRNA arrays including the corresponding SKO crRNA arrays crApc.NTC and crCasp8.NTC (Fig. 3a). Interestingly, this finding that several DKO crRNA arrays were more heavily enriched than their SKO counterparts was corroborated across tumors. For instance, Tumor 3 was dominated by crSetd2.crAcvr2a and crRnf43.crAtrx, Tumor 5 by crCic.crZc3h13 and crCbwd1.crNsd1, and Tumor 6 by crAtm.crRunx1 and crKmt2d.crH2-Q2 (Fig. 3a, Fig. S2d). In all of these cases, the corresponding SKO crRNA arrays were far less abundant compared to the DKO crRNA arrays. Taken together, these data point to the dominance of a handful of individual clones within each tumor sample, and further suggest that certain double-mutant clones had out-competed the corresponding single-mutant clones.

**Figure 3:**
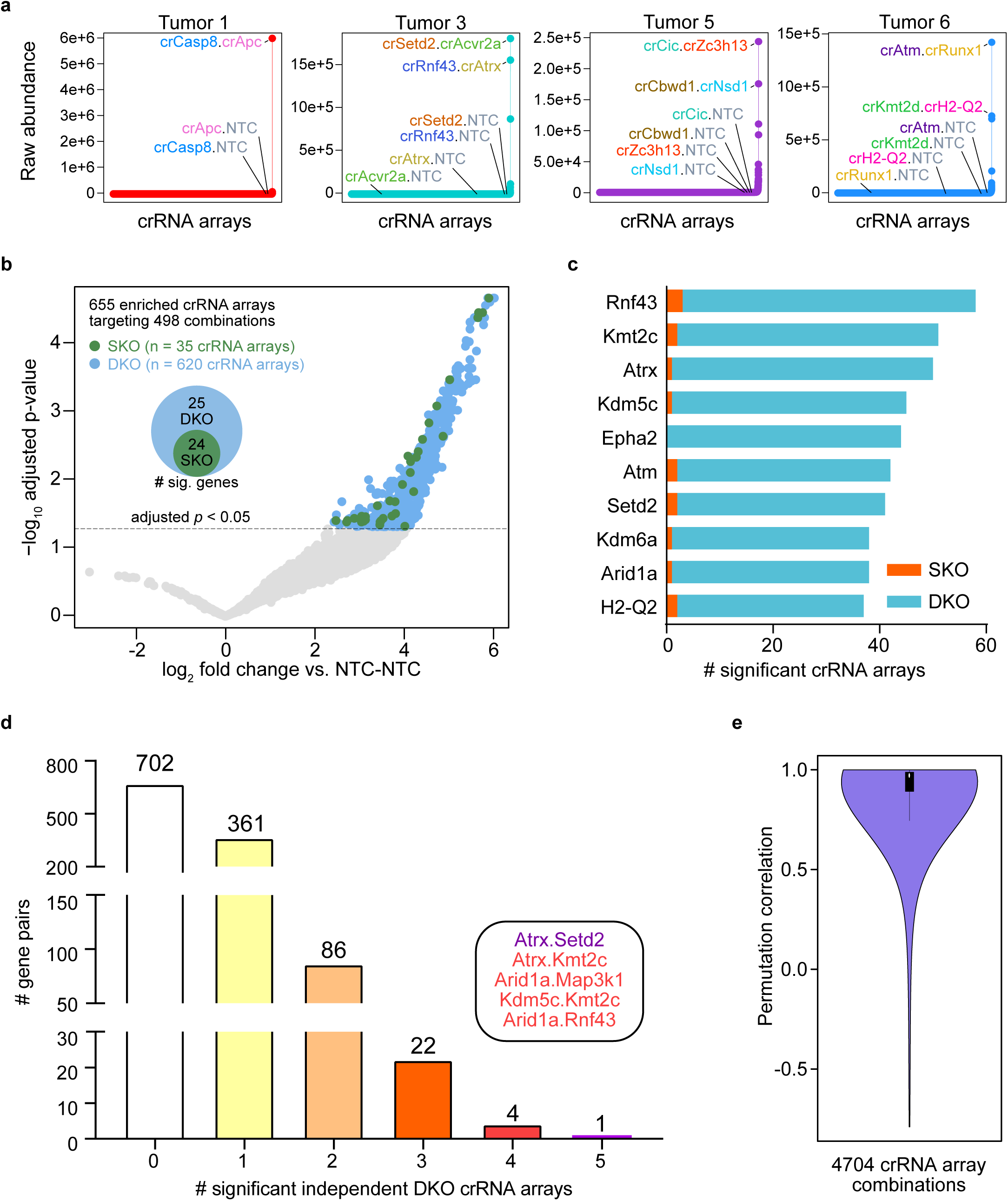
Enrichment analysis of single knockout and double knockout crRNA arrays. **a.** Ranked crRNA array abundance plots of four representative tumor samples. In each tumor, there was a distinct set of DKO crRNA arrays that showed clear enrichment above the rest of the library, including the corresponding SKO crRNA arrays for each DKO pair. In Tumor 1 (red), crCasp8.crApc was by far the most abundant crRNA array, dwarfing all other crRNA arrays including the corresponding SKO crRNA arrays crApc.NTC and crCasp8.NTC. Tumor 3 (teal) was dominated by crSetd2.crAcvr2a and crRnf43.crAtrx, Tumor 5 (purple) by crCic.crZc3h13 and crCbwd1.crNsd1, and Tumor 6 (blue) by crAtm.crRunx1 and crKmt2d.crH2-Q2. In all of these cases, the corresponding SKO crRNA arrays were far less abundant compared to the DKO crRNA arrays. **b.** Volcano plot of DKO (blue) and SKO (green) crRNA arrays compared to NTC-NTC controls. Log2 fold change is calculated using average log2 rpm abundance across all tumor samples, after averaging the 3 NTC-NTC controls to get one NTC-NTC score per sample. 655 crRNA arrays were found to be significantly enriched compared to NTC-NTC controls (Benjamini Hochberg-adjusted p < 0.05). Of these, 620 were DKO crRNA arrays and 354 were SKO crRNA arrays. In total, the 655 enriched crRNA arrays corresponded to 498 gene combinations. A Venn diagram is also shown, detailing the number of genes involved in significant DKO and/or SKO crRNA arrays. All 49 genes in the PANCAN17-mTSG CCAS library were represented within at least one significant DKO crRNA array, while 24 genes were found to be significant as part of a SKO crRNA array. **c.** Bar plot of the top 10 genes ranked by the number of significant crRNA arrays associated with each gene. DKO crRNA array counts are shown in blue, and SKO crRNA arrays in orange. *Rnf43* and *Kmt2c* were the two most influential genes, associated with 58 and 51 independent crRNA arrays. **d.** Bar plot showing the number of significant DKO crRNA arrays associated with each gene pair in the CCAS library. 113 gene pairs were represented by at least 2 independent DKO crRNA arrays. Of note, the interaction of *Atrx*+*Setd2* was supported by 5 independent crRNA arrays, while *Atrx+Kmt2c*, *Arid1a+Map3k1*, *Kdm5c+Kmt2c*, and *Arid1a+Rnf43* were substantiated by 4 crRNA arrays. **e.** Violin plot showing the distribution of permutation correlations between crX.crY and crY.crX for the 4,704 DKO crRNA array combinations in the CCAS library (9,408 unique crRNA array permutations). In total, 80.1% (3,767/4,704) of all crRNA array combinations were significantly correlated when comparing the two permutations associated with each combination (Benjamini-Hochberg adjusted *p* < 0.05, by t-distribution).

In order to uncover the genetic interactions underlying the positive selection *in vivo,* we set out to quantitatively identify all significantly enriched crRNA arrays across all 10 tumors (Methods). We compared the abundance of each DKO and SKO crRNA array to the average of all NTC-NTC crRNA arrays. 655 crRNA arrays targeting 498 gene combinations were found to be significantly enriched compared to NTC-NTC controls (Benjamini-Hochberg adjusted *p* < 0.05) (Fig. 3b; Table S7). Of these, 620 were DKO crRNA arrays and 35 were SKO crRNA arrays. We then decomposed the 655 significantly enriched crRNA arrays to their constituent single crRNAs, and identified the target genes associated with each single crRNA. All 49 genes in the PANCAN17- mTSG CCAS library were represented within at least one significant DKO crRNA array, and 24 genes were additionally found to be significant as part of a SKO crRNA array (Fig. 3b, Table S8). To identify the genes most frequently targeted among the set of 655 significant crRNA arrays, we counted the number of significant crRNA arrays associated with each gene and found that *Rnf43* and *Kmt2c* were the two genes with the largest number of significant crRNA arrays (Fig. 3c, Table S8). Interestingly, of the top 10 genes in this analysis, 6 are epigenetic modifiers (*Kmt2c*, *Atrx, Kdm5c, Setd2, Kdm6a, and Arid1a*), revealing the direct phenotypic consequence of their loss-of-function in tumor suppressor gene networks.

We then sought to identify specific genetic interactions that comprise this network. We quantified the number of significant DKO crRNA arrays associated with each gene pair (Methods) (Fig. 3d, Table S9). 113 gene pairs were represented by at least 2 independent DKO crRNA arrays. Strikingly, the interaction of *Atrx*+*Setd2* was supported by 5 independent crRNA arrays, while *Atrx+Kmt2c*, *Arid1a+Map3k1*, *Kdm5c+Kmt2c*, and *Arid1a+Rnf43* were substantiated by 4 crRNA arrays. In aggregate, these analyses generated an unbiased profile of genetic interactions in tumor suppression dismantled upon Cof1-mediated double-mutagenesis.

Looking to investigate possible positional effects for each individual crRNA in the CCAS library, we directly compared the two permutations of each crRNA array combination (Methods) (Fig. S4a). For each of the 4,704 DKO crRNA array combinations (condensed from 9,408 DKO crRNA array permutations), we calculated the Pearson correlation of crRNA array abundance (i.e. comparing crX.crY to crY.crX) across all tumor samples (subsequently referred to as permutation correlation) (Table S11). Examining the distribution of permutation correlations, we observed a strong skew towards high correlation coefficients (median permutation correlation > 0.97) (Fig. 3e), indicating that for most crRNA array combinations, the positioning of constituent single crRNAs did not affect *in vivo* abundance of the crRNA array (Fig. S4b). In total, 80.1% (3,767/4,704) of all crRNA array combinations were significantly correlated when comparing the two permutations associated with each combination (Benjamini-Hochberg adjusted *p* < 0.05, by t-distribution). The two most significantly correlated crRNA array combinations were between the single crRNAs crH2-Q2.1 and crPten.240, and between crCbwd1.84 and crEpha2.5: crH2-Q2.1_crPten.240 was strongly correlated with the abundance of crPten.240_crH2-Q2.1 across all 10 tumors (R = 0.999, *p* = 2.28 e-19), and a similar trend was observed between crCbwd1.84_crEpha2.5 and crEpha2.5_crCbwd1.84 (R = 0.999, *p* = 7.09 e-19) (Fig. S4c-d).

To quantitate the gross contributions of individual crRNAs to tumorigenesis, we performed marginal distribution meta-analysis of all 98 constituent single crRNAs in the CCAS library (Methods) (Fig. S4e). As the CCAS library was designed with crRNA array orientation as a consideration (Fig. 1c), we calculated the average log2 rpm abundance of all DKO crRNA arrays associated with each single crRNA when present in position 1 or in position 2 of the crRNA array (Table S10). Across all 98 single crRNAs, the average abundance for each single crRNA when in position 1 was significantly correlated with its average abundance when in position 2 (Pearson correlation coefficient (R) = 0.397, *p* = 5.25 e-5 by t-distribution). This finding suggests that a crRNA confers a similar selective advantage regardless of position in the crRNA array considering all other crRNAs it is paired with.

### High-throughput identification of synergistic drivers of transformation and tumorigenesis

To quantitatively investigate the genetic interactions in this model, we then developed a metric of synergy for DKO crRNA arrays. Since the relative abundance of a crRNA array is effectively an estimate of its relative selective advantage *in vivo*, we defined the synergy coefficient (SynCo) for each DKO crRNA array as DKO_xy_ - SKO_x_ - SKO_y_. The DKO_xy_ score is the log_2_ rpm abundance of the DKO crRNA array (i.e., crX.crY) after subtracting average NTC-NTC abundance; SKO_x_ and SKO_y_ scores are defined as the average log_2_ rpm abundance of each SKO crRNA array (3 SKO crRNA arrays associated with each individual crRNA), each after subtracting average NTC-NTC abundance (Fig. 4a). By this definition, a SynCo score >> 0 would indicate that a given DKO crRNA array is synergistic, as the DKO score would thus be greater than the sum of the individual SKO scores on a log-linear scale.

**Figure 4:**
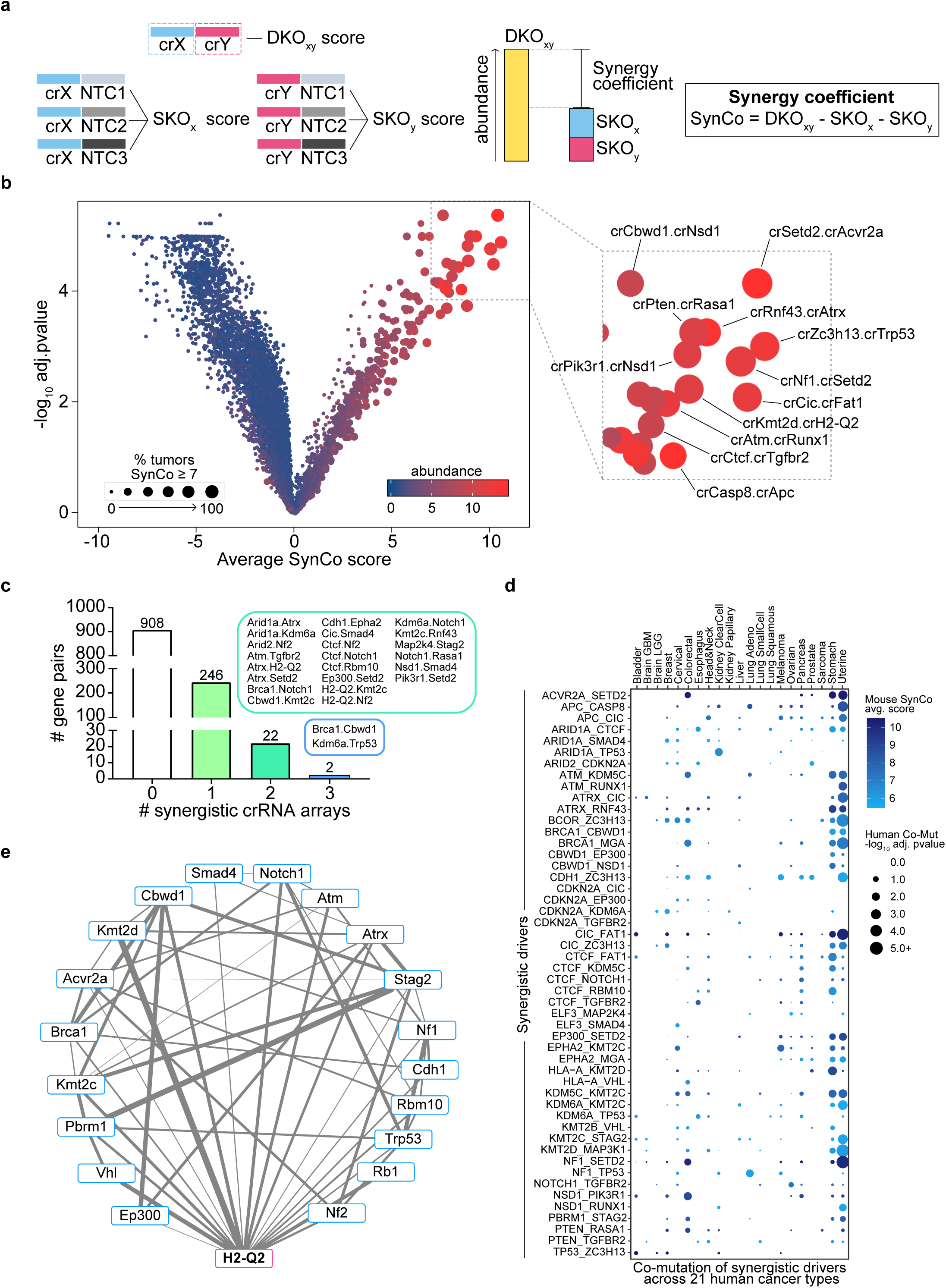
High-throughput identification of synergistic gene pairs as co-drivers of transformation and tumorigenesis. **a**. Schematic describing the methodology for calculating a synergy coefficient (SynCo) for each DKO crRNA array in individual tumor samples. DKO_xy_ score is the log2 rpm abundance of the DKO crRNA array (i.e., crX.crY) after subtracting average NTC-NTC abundance. SKO_x_ and SKO_y_ scores are defined as the average log2 rpm abundance of each SKO crRNA array (3 SKO crRNA arrays associated with each individual crRNA), after subtracting average NTC-NTC abundance. SynCo = DKOxy - SKO_x_ - SKO_y_. By this definition, a SynCo score >> 0 would indicate that a given DKO crRNA array is synergistic, as the DKO score would thus be greater than the sum of the individual SKO scores. **b.** Volcano plot of average SynCo across all tumor samples and associated -log_10_ Benjamini-Hochberg adjusted p-value (two-sided one sample t-test, H_0_: mean SynCo = 0) for each DKO crRNA array in the library. Each point is scaled by size, in reference to the % of tumor samples with a SynCo ≥ 7 for a particular crRNA array, and also color-coded according to the average log2 rpm abundance across all tumor samples. To the right is a zoomed-in view of the top synergistic DKO crRNA arrays. Among the strongest driver pairs were crSetd2.crAcvr2a, crCbwd1.crNsd1, crRnf43.crAtrx, and crPten.crRasa1. **c.** Bar plot showing the number of significantly synergistic dual-crRNAs associated with each gene pair in the CCAS library (Benjamini-Hochberg adjusted *p* < 0.05). 24 synergistic pairs were corroborated by multiple dualcrRNAs, including *Brca1*+*Cbwd1* and *Kdm6a+Trp53*. **d.** Gene-level synergistic driver network based on the CCAS screen, focusing here on *H2-Q2* and all first-degree connections between genes associated with *H2-Q2*. The complete network is shown in **Fig. S5**. Each node represents one gene, and each edge indicates a significant synergistic interaction (Benjamini-Hochberg adjusted *p* < 0.05). Edge widths are scaled by SynCo score. *H2-Q2* was significantly synergistic with a total of 19 other genes by this analysis, and its strongest synergistic partner was found to be *Kmt2d* (SynCo = 8.877). **e.** Bubble chart depicting co-mutation analysis of synergistic drivers across 21 human cancer types. For each of the top 50 significant driver pairs identified through CCAS SynCo analysis, bubble dots indicate whether these gene pairs were significantly co-mutated in human cancers (where mutations are defined as nonsynonymous mutations or deep deletions). The color of each point corresponds to the average SynCo score (from mice), while the size of each point is scaled to the -log_10_ p-value of co-mutation in each human cancer (hypergeometric test). Of all synergistic interactions identified by SynCo analysis, 132 gene combinations were significantly co-mutated in at least one cancer type (Benjamini-Hochberg adjusted *p* < 0.05), with 46 pairs significantly co-mutated in two or more cancer types, indicating that the synergistic driver pairs identified through the mouse CCAS screen recapitulate genomic features of human cancers.

We calculated the SynCo of each DKO crRNA array within each tumor sample (Table S12), and assessed whether the SynCo score of a given crRNA array across all 10 tumors was statistically significantly different from 0 by a two-sided one-sample t-test (Table S13). Out of 9,408 DKO crRNA arrays, 294 were significantly synergistic (Benjamini-Hochberg adjusted *p* < 0.05, average Synco > 0), representing 270 gene combinations. To obtain a comprehensive picture of the synergistic driver pairs, we plotted the average SynCo of each DKO crRNA array against its associated p-value, while additionally color-coding each point by average abundance and scaling the size of each point by the percentage of tumors that had a high SynCo score (SynCo ≥ 7) for that crRNA array (Fig. 4b). Among the top synergistic driver pairs in this analysis were crSetd2.crAcvr2a and crCbwd1.crNsd1. *Setd2* encodes a histone methyltransferase that has been implicated in a number of cancer types ^28–30^, while *Acvr2a* is a receptor serine-threonine kinase that plays a critical role in Tgf-β signaling and is frequently mutated in microsatellite-unstable colon cancers ^31,32^. *Nsd1* encodes a lysine histone methyltransferase that has been linked to Sotos syndrome, a genetic disorder of cerebral gigantism, and has been implicated in various cancers ^33–35^. In contrast, *Cbwd1* encodes an evolutionarily conserved protein whose biological function is unknown; on the basis of its amino acid sequence, *Cbwd1* has been predicted to contain a cobalamin synthase W domain ^36^, but its function has never been characterized in a mammalian species. Interestingly, many of the high-score SynCo-significant gene pairs have not been functionally characterized in literature.

To pinpoint the most robust genetic interactions from SynCo analysis, we quantified the number of synergistic dual-crRNAs associated with each gene pair. Of the 268 significant gene pairs, 24 were represented by at least 2 synergistic dual-crRNAs (Fig. 4c). Considering many gene pairs might have additive effects, the SynCo score is a stringent metric of genetic interaction; thus the finding that several gene pairs were further supported by multiple synergistic dual-crRNAs provides further evidence for the genetic interactions between these genes.

Having identified 270 significant pairwise genetic interactions in early tumorigenesis, many of which corresponded to genomic features of human tumors, we next sought to place each of these gene pairs within the larger network of tumor suppression. We constructed a network of all synergistic driver interactions captured by CCAS screening, where each node represents a gene and each edge represents a significant synergistic interaction (Fig. S5). In this network, the color of each gene is scaled by its degree of connectivity, while edge widths are scaled to the SynCo score associated with that interaction. Surprisingly, *H2-Q2,* a gene encoding a major histocompatibility complex (MHC) component, the murine homolog of human HLA-A MHC class I A, was found to have the greatest network connectivity, with 19 different interacting partners (Fig. 4d). Of note, *H2-Q2* shared its strongest interaction with *Kmt2d* (SynCo = 8.877), pointing to a genetic interaction between an epigenetic modifier and an immune regulator in tumorigenesis. Many of these synergistic pairs were significantly co-mutated in one or more cancer types (top 50 SynCo interactions shown in Fig. 4e), suggesting relevance of these genomic features in human cancers.

### Cpf1 crRNA array library screen in a mouse model of metastasis

To further demonstrate the versatility, we performed Cpf1 crRNA array library screening in a mouse model of metastasis to identify co-drivers of metastatic process *in vivo*. We generated lentiviral pools from the CCAS plasmid library, and subsequently infected Cpf1+ KPD cells to perform massively parallel gene-pair level mutagenesis. We then injected the mixed double mutant cell populations (CCAS-treated cells, 4×10^6^ cells per mouse, ~400x coverage) subcutaneously into *Nu/Nu* mice (n = 7) and *Rag*1-/- mice (n = 4). After 8 weeks, we collected the primary tumors, four lung lobes, and other stereoscope-visible metastases (two large extra-pulmonary metastases were found), and then subjected to crRNA array sequencing (Fig. 5a). The 3 pre-injection cell pools, as well as primary tumors and metastases from all 11 mice were sequenced (Methods) (Table S16). As seen in the overall representation of the CCAS library across all metastasis screen samples (Fig. 5b, Fig. S7) (Tables S17, S18), whereas cell samples exhibited lognormal representation of the CCAS library, both primary tumors and metastases showed strong enrichment of specific SKO and DKO crRNA arrays. NTC-NTC crRNA arrays were consistently found at low abundance in all primary tumors and metastases samples, indicating strong selection and clonal expansion during the metastasis process. Notably, the crRNA library representation of metastases in all the collected lobes showed high degree of similarity to primary tumors (Fig. 5c), consistent with a common clonal origin from the same primary tumors within each individual mouse.

**Figure 5:**
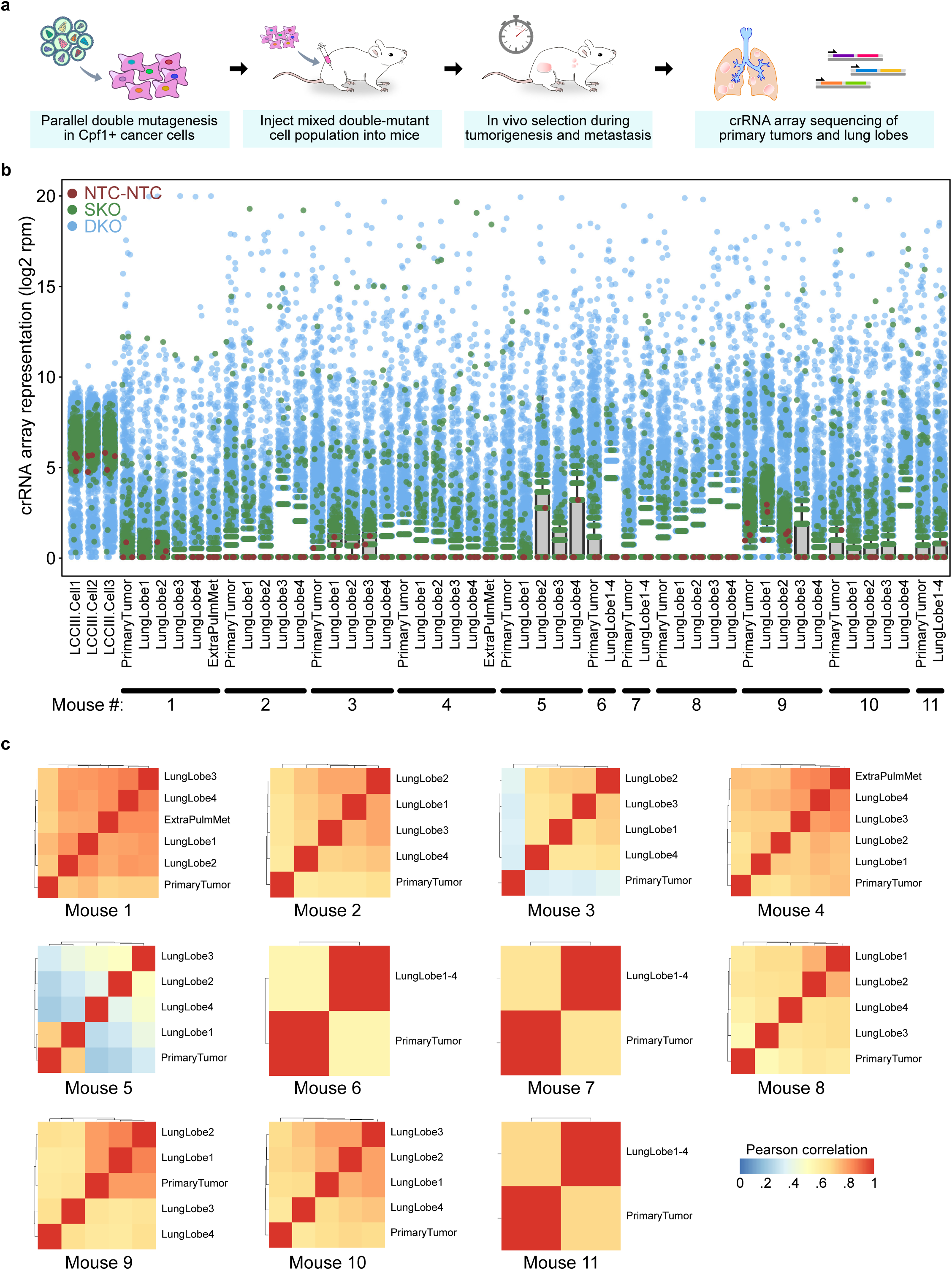
Cpf1 crRNA array library screen in a mouse model of metastasis. **a.** Schematic of the experimental approach for Cpf1 crRNA array library screen in a mouse model of metastasis to identify co-drivers of metastatic process *in vivo*. We generated lentiviral pools from the CCAS plasmid library, and subsequently infected Cpf1+ KPD LCC cells to perform massively parallel gene-pair level mutagenesis. The mixed double mutant cell populations (CCAS-treated cells, 4× 10^6^ cells per mouse, ~400x coverage) were then injected subcutaneously into *Nu/Nu* mice (n = 7) and *Rag1*-/- mice (n = 4). After 8 weeks, genomic DNA was extracted from the primary tumors, four lung lobes, and other stereoscope-visible metastases, and then subjected to crRNA array sequencing. **b.** Dot-boxplot depicting the overall representation of the CCAS library across all metastasis screen samples, in terms of log2 rpm abundance. The 3 pre-injection cell pools, as well as primary tumors and metastases from all 11 mice were sequenced. NTC-NTC controls are shown in red, SKO crRNA arrays in green, and DKO crRNA arrays in blue. Whereas cell samples exhibited lognormal representation of the CCAS library, both primary tumors and metastases showed strong enrichment of specific SKO and DKO crRNA arrays. Notably, NTC-NTC crRNA arrays were consistently found at low abundance in all primary tumors and metastases samples. **c.** Intra-mouse Pearson correlation heatmaps of samples, showing high degree of similarity between primary tumors and metastases from the same host.

### Enrichment analysis of crRNA arrays identified metastasis drivers and co-drivers

In the CCAS metastasis screen dataset, we again observed strong overall permutation correlation, where 97.4% all crRNA array combinations were significantly correlated when comparing the two permutations associated with each combination (Benjamini-Hochberg adjusted *p* < 0.05, by t-distribution) (median permutation correlation > 0.85) (Fig. 6a), indicating that for most crRNA array combinations, the positioning of constituent single crRNAs did not affect *in vivo* abundance of the crRNA array. We then compared DKO and SKO crRNA arrays to NTC-NTC controls in the metastasis screen. Across all *in vivo* samples, 2933 crRNA arrays were found to be significantly enriched compared to NTC-NTC controls (Benjamini Hochberg-adjusted p < 0.05), targeting 1006 combinations. Of these, 2813 were DKO crRNA arrays and 121 were SKO crRNA arrays (Fig. 6b) (Table S19). All 49 genes in the PANCAN17-mTSG CCAS library were represented within at least one significant DKO crRNA array. The top 15 genes associated with these 2933 crRNA arrays ranked by the number of significant crRNA arrays associated with each gene were found to be *Arid1a, Cdh1, Kdm5c, Rb1, Epha2, Kmt2b, Cic, Kmt2c, Kdm6a, Atra, Nf2, Elf3, Apc, Rnf43* and *Ctcf* (Fig. 6c) (Table S20).

**Figure 6:**
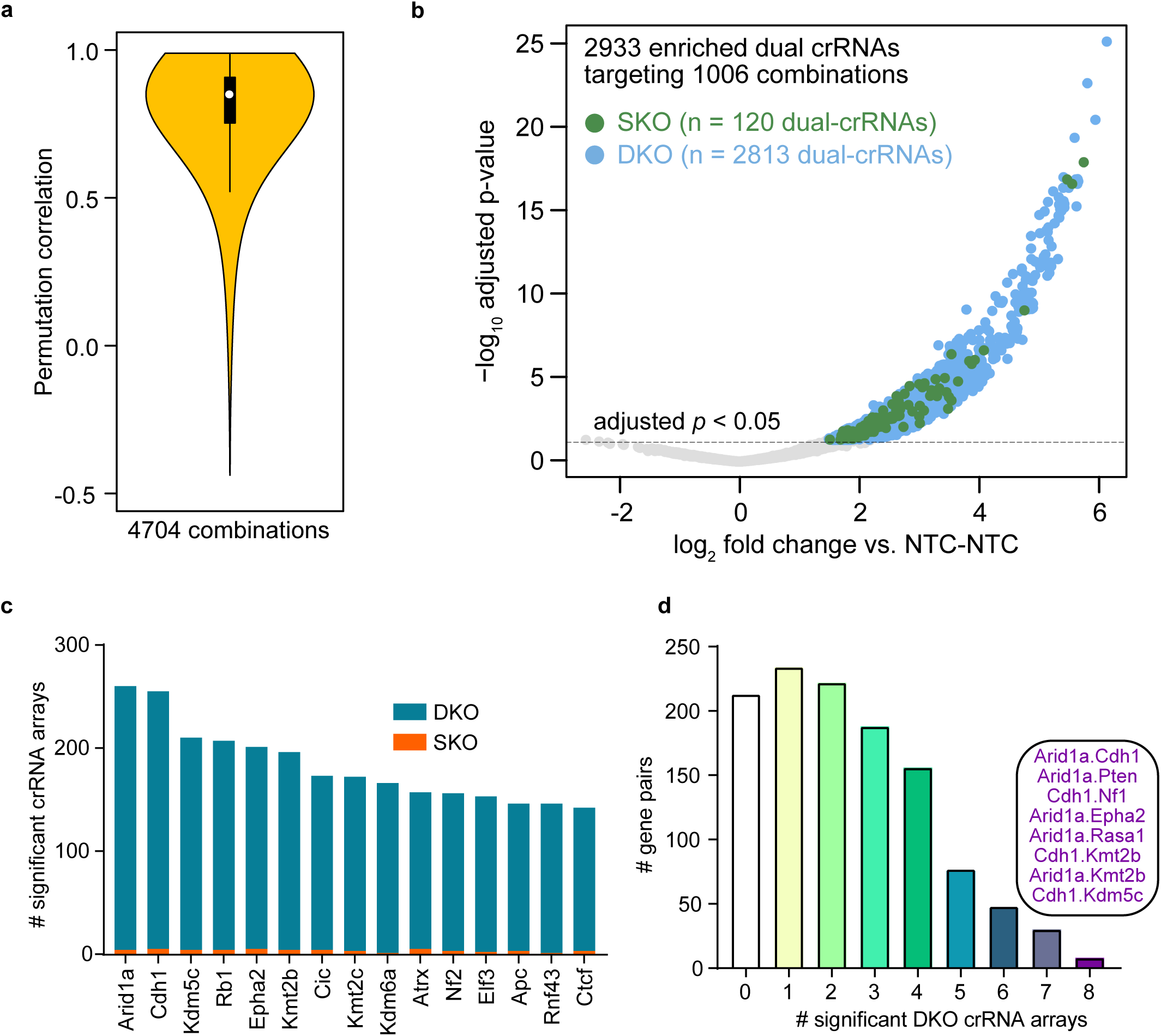
Enrichment analysis of crRNA arrays identified metastasis drivers and co-drivers. **a.** Violin plot showing the distribution of permutation correlations between crX.crY and crY.crX for the 4,704 DKO crRNA array combinations in the CCAS library (9,408 unique crRNA array permutations). 97.4% all crRNA array combinations were significantly correlated when comparing the two permutations associated with each combination (Benjamini-Hochberg adjusted *p* < 0.05, by t-distribution). **b.** Volcano plot of DKO (blue) and SKO (green) crRNA arrays compared to NTC-NTC controls in the metastasis screen. Log2 fold change is calculated using average log_2_ rpm abundance across all *in vivo* samples, after averaging the 3 NTC-NTC controls to get one NTC-NTC score per sample. 2933 crRNA arrays were found to be significantly enriched compared to NTC-NTC controls (Benjamini Hochberg-adjusted p < 0.05), targeting 1006 gene pairs. Of these, 2813 were DKO crRNA arrays and 120 were SKO crRNA arrays. All 49 genes in the PANCAN17-mTSG CCAS library were represented within at least one significant DKO crRNA array. **c.** Bar plot of the top 15 genes ranked by the number of significant crRNA arrays associated with each gene. DKO crRNA array counts are shown in blue, and SKO crRNA arrays in orange. *Arid1a, Cdh1, Kdm5c* and *Rb1* were the top genes associated with ≥ 200 independent crRNA arrays. **d.** Bar plot showing the number of significant DKO crRNA arrays associated with each gene pair in the CCAS library. Most gene pairs were represented by at least 2 independent DKO crRNA arrays. Of note, 8 gene pairs were represented by all eight crRNA arrays.

We then sought independent evidence for selection of metastasis co-drivers via investigation of independent crRNA arrays targeting the same gene pair. By calculating the number of significant DKO crRNA arrays associated with each gene pair in the CCAS library, we found that the majority (729/1176 = 61.99%) of gene pairs were represented by at least 2 independent DKO crRNA arrays (Table S21). Of note, 30 gene pairs were represented by seven independent crRNA arrays, among them including *Apc+Cdh1, Cdh1+H2-Q2, Epha2+ Kmt2b*; and 8 gene pairs were represented by all eight designed crRNA arrays, including *Arid1 a+Pten, Cdh1+Nf1, Cdh1+Kdm5c, Arid1a+Rasa1, Arid1a+Cdh1 Cdh1+Kmt2b, Arid1a+Kmt2b, and Arid1a+Epha2,* suggesting these are the strong co-drivers of metastasis (Fig. 6d, Table S21).

### Modes and patterns of metastatic spread with co-drivers

We then investigated the *in vivo* patterns of metastatic evolution of these double mutants. Examination of the clonal architecture of the crRNA arrays in the metastases samples revealed a highly heterogenous pattern of clonal dominance (Fig. S8). Comparison of the crRNA array representations between metastases to primary tumors revealed modes of monoclonal spread (Fig. 7a, Fig. S8) where dominant metastases in multiple lobes were derived from identical crRNA arrays, as well as polyclonal spread (Fig. 7b, Fig. S8) where dominant metastases in all lobes were derived from multiple varying crRNAs. For example, mouse 1 represents a case of monoclonal spread where all 4 lobes were dominated by a clone, crNf2.crRnf43, that was also found at the primary tumor as a major clone (>=2% frequency). In contrast, mouse 10 represents a case of polyclonal spread where each lung lobe were comprised of a myriad of crRNA arrays. Namely, lobes 1 and 2 were dominated by crNsd1.crNTC, and crH2-Q2.crCdh1 + crNsd1.crAtm + crCasp8.crArid1a, respectively, which were also major clones in primary tumor (Fig. 7b). However, lobe 3 was dominated by crElf3.crFbxw7 + crRb1.crCasp8, which were not found as major clones in primary tumor; the case of lobe 4 echoes that of lobe 3 with a more complex metastatic clonal mixture, in which most of the dominant clones (crBcor.crKdm5c, crAcvr2a.crNTC, crRb1.crCasp8, crCdkn2a.crApc, crApc.crKmt2b, crRasa1.crNf2, crElf3.crFbxw7 and crPten.crKdm5c) were not found as major clones in the primary tumor (Fig. 7b).

**Figure 7:**
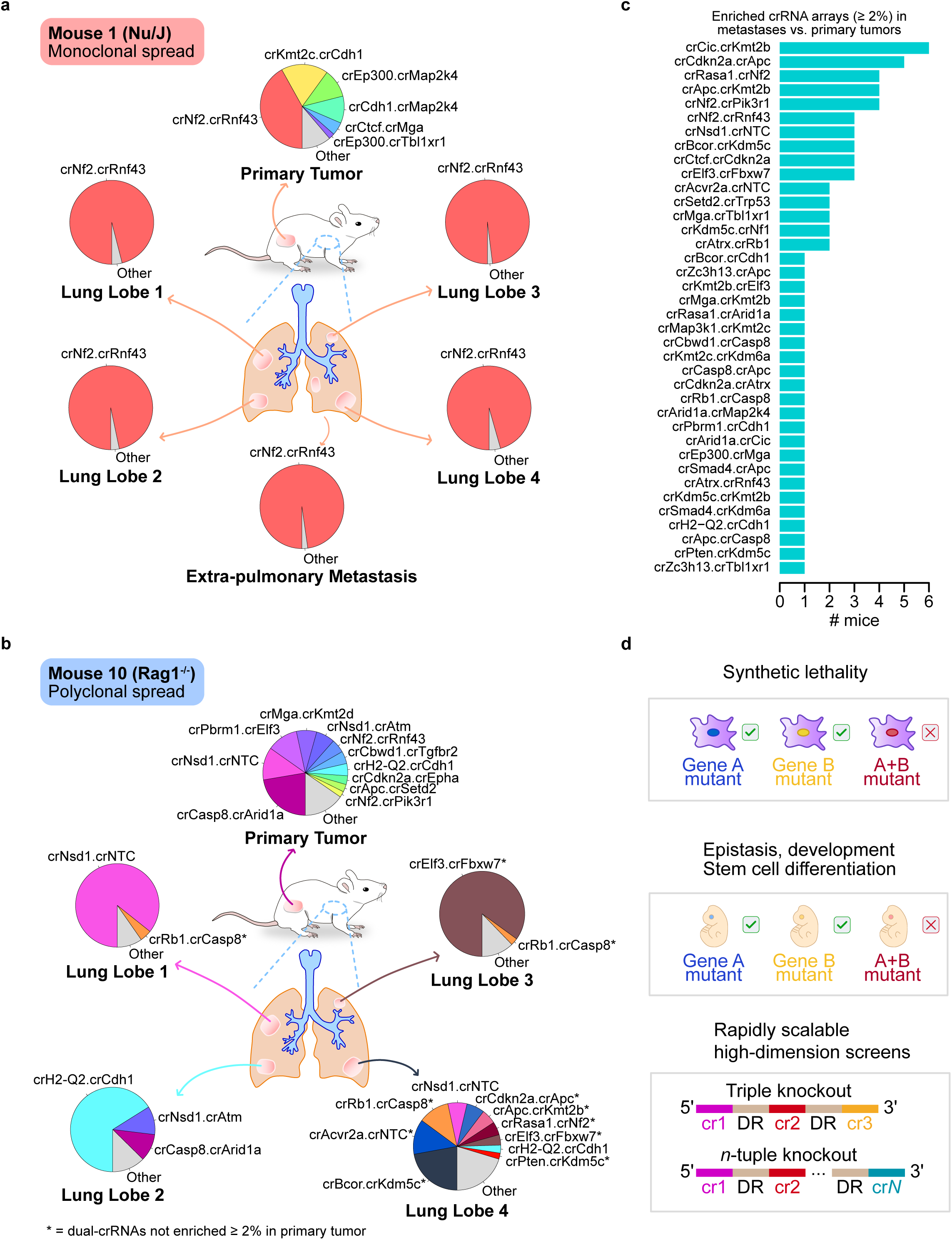
Modes and patterns of metastatic spread with co-drivers. Comparison of the crRNA array representations between metastases to primary tumors revealed modes of monoclonal spread (A) where dominant metastases in all lobes were derived from identical crRNA arrays, and polyclonal spread (B) where dominant metastases in all lobes were derived from several different crRNAs. **a.** Example of a monoclonal spread where all 4 lobes were dominated by a clone crNf2.crRnf43, that was also found at the primary tumor as a major clone (≥ 2% frequency). **b.** Example of a polyclonal spread where all 4 lobes were derived from multiple varying crRNAs. Lobe 1 was dominated by crNsd1.crNTC, which was one of major clones in the corresponding primary tumor; Lobe 2 was dominated by crH2-Q2.crCdh1, crNsd1.crAtm and crCasp8.crArid1a, which were also major clones in primary tumor. However, lobe 3 was dominated by crElf3.crFbxw7 and crRb1.crCasp8, which were not found as major clones in primary tumor; the case of lobe 4 echoes that of lobe 3 with a more complex metastatic clonal mixture, as most of its dominant clones (crBcor.crKdm5c, crAcvr2a.crNTC, crRb1.crCasp8, crCdkn2a.crApc, crApc.crKmt2b, crRasa1.crNf2, crElf3.crFbxw7 and crPten.crKdm5c) were not found as major clones in the primary tumor. **c.** Waterfall plot of enriched crRNA arrays in a metastases vs primary tumor analysis, identifying crRNA arrays that were dominant clones in metastases but not in the corresponding primary tumor. Top ranked metastasis-specific dominant crRNA arrays were found to be crCic.crKmt2b, crCdkn2a.crApc, crRasa1.crNf2, crApc.crKmt2b, crNf2.crPik3r1, crNf2.crRnf43, among 23 enriched crRNA arrays. **d.** Schematic describing several extended applications of multiplexed Cpf1 screens. The relative ease of library construction and subsequent readout with our approach empowers the study of previously intractable biological problems, including combinatorial genome-wide knockout studies of synthetic lethality, as well as the discovery and characterization of epistatic networks in embryonic development and stem cell differentiation. Notably, this approach is rapidly scalable to triple knockout or higher-dimensional screens.

To quantify the metastasis-specific signature of double mutants, we calculated the number of times a crRNA array was considered as metastasis-enriched (i.e. a dominant clone in a lung lobe or extra-pulmonary metastasis (>=2% total reads) but not a dominant clone in the corresponding primary tumor of the same mouse). Top ranked metastasis-specific dominant crRNA arrays were found to be crCic.crKmt2b, crCdkn2a.crApc, crRasa1.crNf2, crApc.crKmt2b, crNf2.crPik3r1, crNf2.crRnf43, among 23 enriched crRNA arrays, with crCic.crKmt2b being metastasis-enriched 55% (6/11 mice) of the time. These data suggest strong genetic signatures of metastasis-specific co-drivers, which have notably been difficult to parse from single-gene studies. Collectively, our results demonstrate the power of *in vivo* Cpf1 crRNA array screens for mapping and identification of genetic interactions in an unbiased manner.

## Discussion

Due to the complex nature of biological systems, a single gene is often far from sufficient to explain the biological or pathological variation observed in health and disease ^2,3,12,14^. Genetic interactions are the building blocks of highly connected biological networks, and their modular nature enables biological pathways to take on a variety of forms – linear, branching divergent, convergent, feed-forward, feedback, or any combination of the above ^3,37,38^. In systems biology, numerous theories and algorithms have been developed to understand such complex networks and to predict genetic interactions. However, predictions have often been surprised by unexpected experimental findings, urging for experimental testing of combinatorial perturbations in a systems manner ^5^.

High-throughput genetic screens are a powerful approach for mapping genes to their associated phenotypes ^39,40^. Unbiased and quantitative analysis of double knockouts enables phenotypic assessment of all possible combinations of any given gene pairs ^41, 42^. Advances in high-throughput technologies utilizing RNA-interference-based gene knockdown ^39^ or CRISPR/Cas9-based gene knockout, activation and repression ^43–45^, have enabled genome-scale screening in multiple species across various biological applications ^46–48^. While high-throughput genetic perturbation approaches have been developed to map out the landscape of genetic interactions in yeast ^49–53^ and in worms ^54,55^, large-scale double knockout studies in mammalian species are scarce, due to the exponentially scaling number of possible gene combinations and the technological challenges of generating and screening double knockouts. Recently, several high-throughput double perturbations have been performed in mammalian cells using RNA interference (RNAi) or clustered regularly interspaced short palindromic repeats (CRISPR)/Cas9 technologies ^56–60^. However, RNAi-based methods act on the level of mRNA silencing. Though CRISPR/Cas9-based methods can induce complete knockouts, the dependence of Cas9 on a trans-activating crRNA (tracrRNA) requires multiple sgRNA cassettes, hindering the scalability of Cas9-mediated high-dimensional screens, and making *in vivo* genetics more difficult.

Cpf1 was recently identified and characterized as a single-effector RNA-guided endonuclease with two orthologs from Acidaminococcus (AsCpf1) and Lachnospiraceae (LbCpf1) capable of efficient genome-editing activity in human cells ^15–19,61^. Unlike Cas9, Cpf1 requires only a single 39-42-nt crRNA without the need of an additional trans-activating crRNA, enabling one RNA polymerase III promoter to drive an array of several crRNAs targeting multiple loci simultaneously ^62,63,64^. This unique feature of the Cpf1 nuclease greatly simplifies the design, synthesis and readout of multiplexed CRISPR screens, making it a suitable system to carry out combinatorial screens. Recognizing the potential of Cpf1 for genome editing applications, rapid progress has been made with several studies focused on Cpf1, including genome-wide measurement of off-target Cpf1 activity ^61,65^, embryonic genome editing in mice ^66,67^, and solution of Cpf1 crystal structures ^63,64^.

Considering that cancer is a polygenic disease of malignant somatic cells, we first designed and performed a Cpf1 double knockout screen in a mouse model of malignant transformation and early tumorigenesis. In this setting, we demonstrated successful mapping of all permutations of crRNA arrays targeting combinations of two putative non-oncogenes, revealing a wide array of unexpected synergistic gene pairs. We found that the most highly connected ‘hub’ genes were epigenetic factors such as *Kmt2c, Atrx, Kdm5c, Setd2, Kdm6a,* and *Arid1a*, suggesting that the multifarious interactions of these factors, whether direct or indirect, lead to drastically accelerated tumorigenesis upon loss-of-function. This finding might explain why, despite being frequently mutated in human cancers ^26^, single knockouts of such factors rarely lead to tumorigenesis *in vivo* (though only a limited number of these genes have thus far been studied in animal models) ^29, 68^. In that sense, epigenetic modifiers might function as genetic buffers, redundant backup pathways, modifiers or amplifiers of multiple other apparently unrelated pathways ^55^. Many of the synergistic interactions identified through our screen were subsequently found to be significantly co-mutated across multiple cancer types. In a more complex biological process such as metastasis, which includes a cascade of primary tumor growth, inducing angiogenesis and lyphangiogenesis, extravasation, circulation, extravasation, colonization and immunological interactions, our screen is capable of detecting robust signatures of selection and revealing modes and patterns of clonal expansion of complex pools of double mutants *in vivo*. Multiplexed Cpf1 screens thus represent a powerful tool for studying genetic interactions with unparalleled simplicity and specificity.

We showed here that multiplexed Cpf1 screens can enable the high-throughput discovery of synergistic interactions by examining patterns of crRNA array enrichment; on the flip side, crRNA array depletion screens would enable the identification of synthetically lethal gene mutations in cancer, potentially opening new avenues for therapeutic discovery (Fig. 7d). While we focused on TSGs in the present study, CCAS screens can be easily tailored for any particular gene set in any biological context. The present study serves as a proof-of-principle with an unbiased, medium size library targeting all pairwise combinations of a selected set of genes. More comprehensive combinatorial screens will be feasible through this approach, simply by increasing the number and complexity of crRNA arrays in the library, as well as expanding the target cell pool and/or number of experimental animals accordingly. Considering that Cpf1 can easily target more than two loci with a single crRNA array, in the future, multiplexing 3 or more crRNAs in each array will enable direct screens of triple knockouts and even higher-dimension genetic interactions *in vivo*.

## Supplementary Figure Legends

**Supplementary Figure S1:**
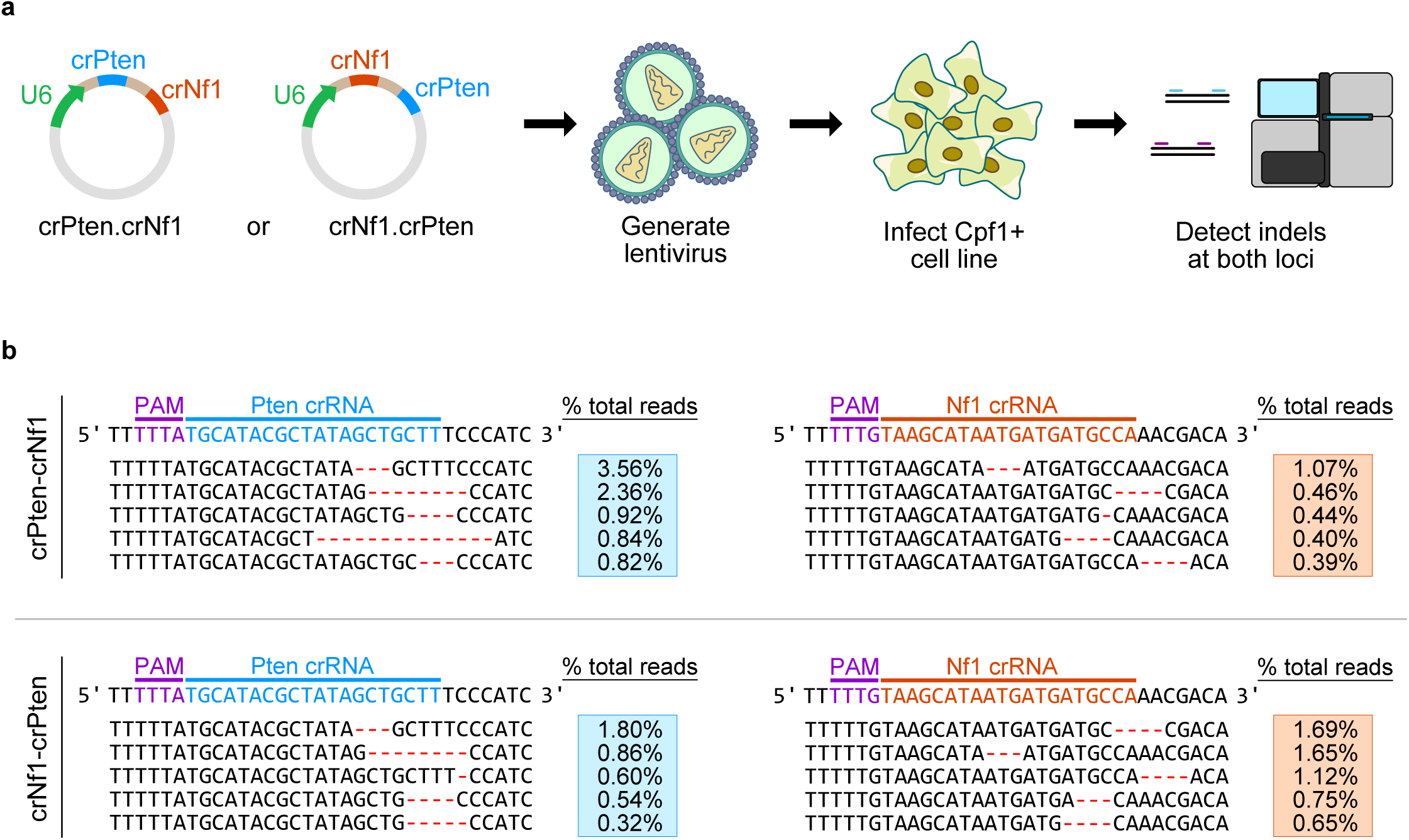
Double knockout of *Nf1* and *Pten* by a single crRNA array. **a.** Schematic depicting the experimental approach for testing the ability of a single crRNA array to induce mutagenesis at both *Nf1* and *Pten*. Plasmids were designed containing a U6 promoter driving the expression of either a *Pten* crRNA (crPten) followed by an *Nf1* crRNA (crNf1), or vice versa. Lentiviruses were subsequently generated and used to infect a tumor cell line that had been transduced with a Cpf1 expression vector (KPD.LbCpf1+). **b.** 7 days after lentiviral infection, genomic DNA was harvested from puromycin-resistant cells for mutation analysis. Nextera library preparation and deep sequencing enabled quantitative high-resolution analysis of the mutations induced by Cpf1 activity. For each treatment condition, mutations were identified at the genomic loci targeted by crPten (left column, blue) and by crNf1) right column, orange). Variant frequencies associated with each mutation are shown in the blue boxes to the right; for each condition, the top 5 most frequent variants are shown. The location of the protospacer adjacent motif (purple) and the crRNA are indicated at the top. Regardless of individual crRNA position within the crRNA array (top row, crPten-crNf1; bottom row, crNf1-crPten), indels were found at both *Pten* and *Nf1* loci in KPD.LbCpf1+ cells treated with crPten-crNf1 or crNf1-crPten crRNA arrays.

**Supplementary Figure S2:**
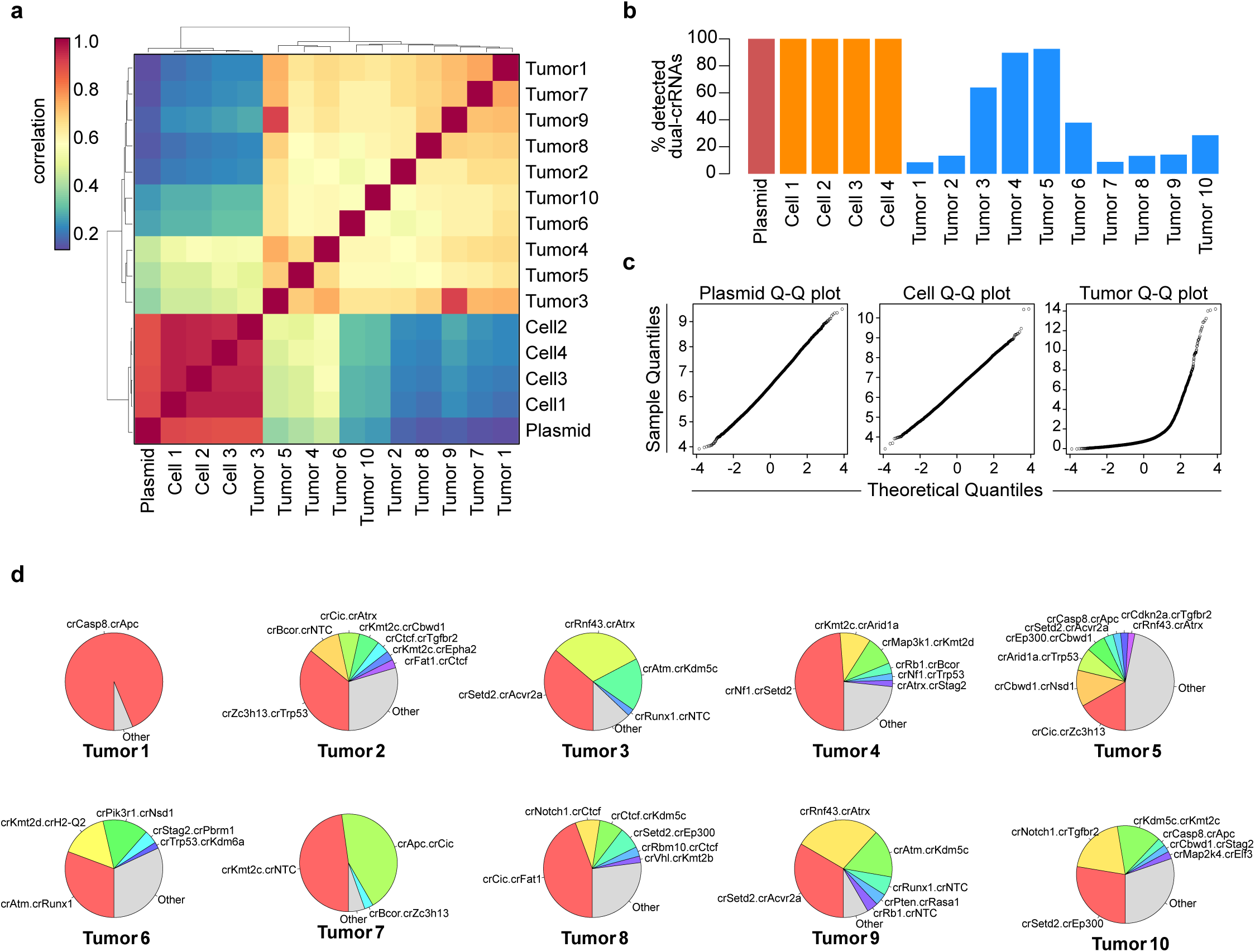
Representation of CCAS crRNA array library in plasmid, cells, and tumors. **a.** Heatmap of pairwise Pearson correlation coefficients of crRNA array log_2_ rpm abundance from CCAS plasmid library, CCAS transduced cells before transplantation (day 7 post infection), and late stage subcutaneous tumors (6.5 weeks post transplantation). Plasmid and cell samples were highly correlated with one another, while tumor samples were most correlated with other tumors. **b.** Bar plot depicting the percentage of all crRNA arrays in the CCAS library that were detected in each sample. All plasmid and cell samples contained 100% of CCAS crRNA arrays, while tumor samples exhibited significantly lower crRNA array library diversity (mean ± SEM = 37.0% ±10.5%; *p =* 2.02 e-4 compared to plasmid and cells, t-test). **c.** Q-Q plots comparing theoretical and sample quantiles of log2 rpm crRNA array abundance in plasmid, cell, and tumor samples (cells and tumor samples averaged by group). In contrast with plasmid and cell samples, tumor samples did not appear linear on the Q-Q plot, indicating that the distribution of crRNA array abundance in plasmid and cell samples (but not tumor samples) approximated a normal distribution. **d.** Pie charts showing highly enriched crRNA arrays (>2% reads) across all 10 tumors; the area for each crRNA array corresponds to the percentage of reads within the tumor.

**Supplementary Figure S3:**
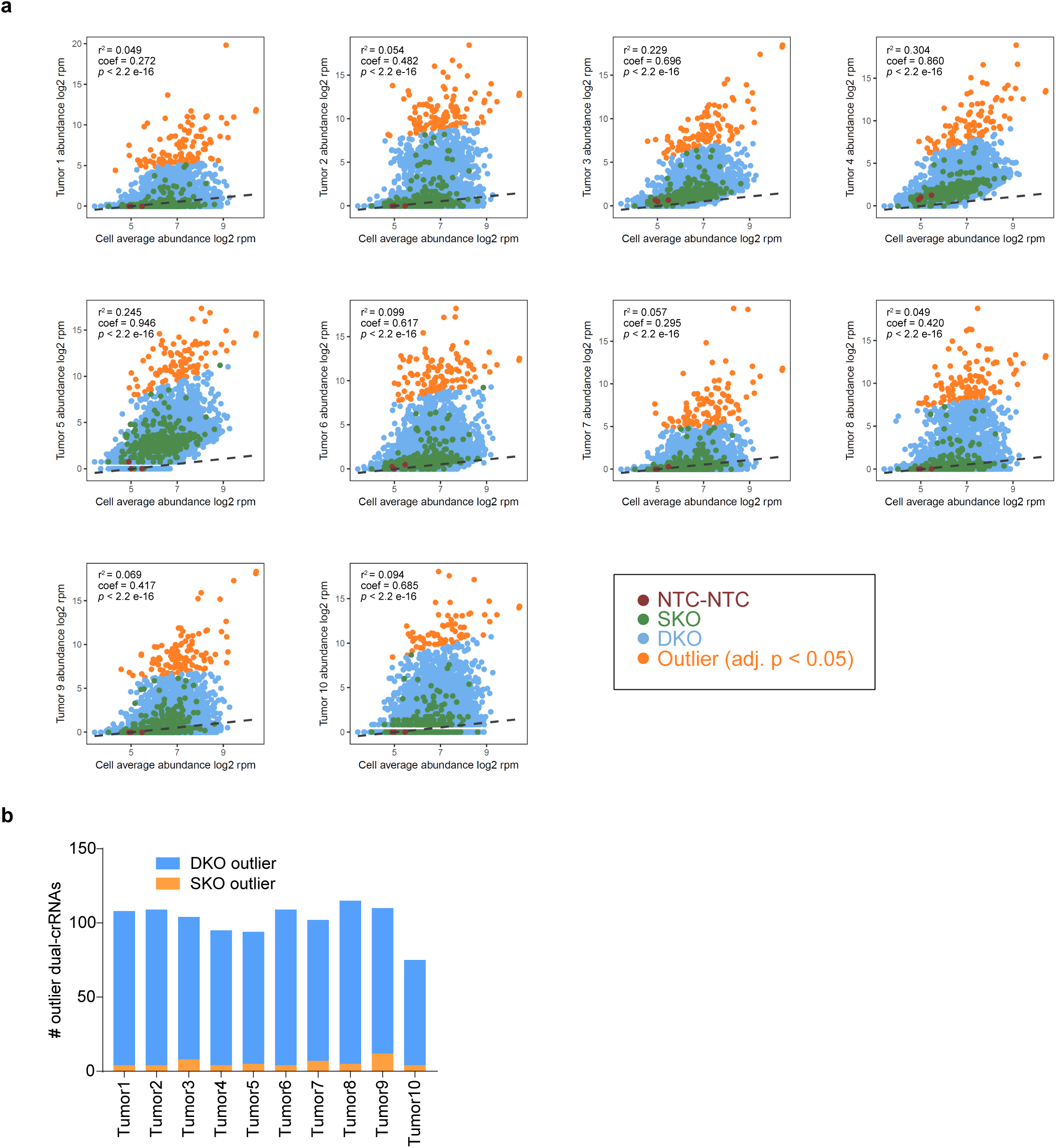
Outlier analysis of individual tumors compared to cells. **a.** Scatterplots comparing log_2_ rpm abundance of crRNA arrays in individual tumors compared to cell samples (cell samples were averaged). In all tumors, crRNA arrays largely approximated a log-linear distribution, as indicated by the linear regression lines (blue = DKO, green = SKO). However, there were numerous clear outliers (orange) (Bonferroni adjusted *p* < 0.05), indicating that specific crRNA arrays had undergone positive selection *in vivo*. The associated regression r^2^, coefficient, and *p-*value (by F-test) are noted on each plot. **b.** Barplots depicting the number of DKO and SKO outlier crRNA arrays identified within each individual tumor, as defined in **a**.

**Supplementary Figure S4:**
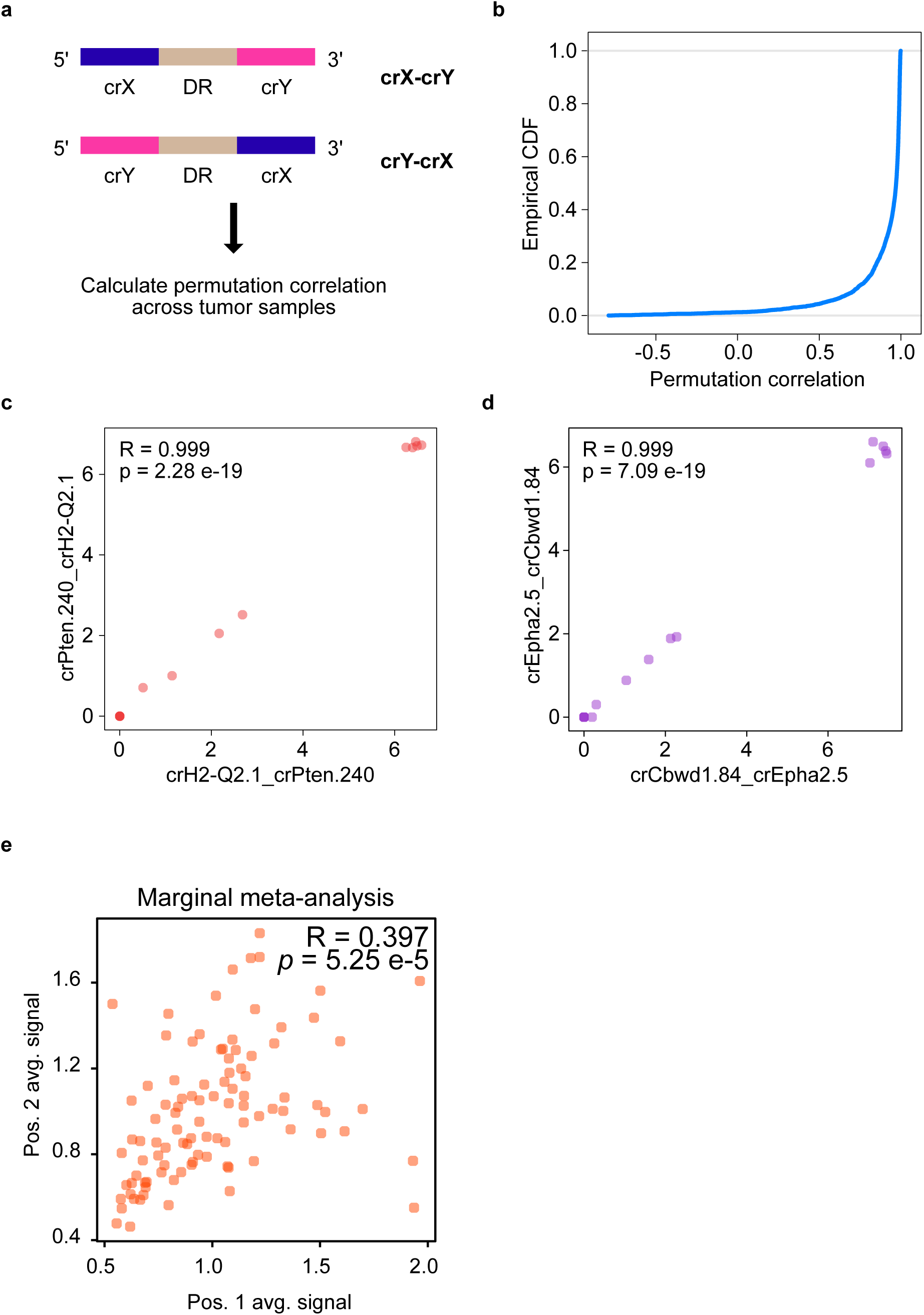
CrRNA array permutation has a minimal effect on enrichment. **a.** Schematic illustrating two permutations of the same crRNA array combination (crX-crY and crY-crX). To estimate possible position effects on the efficiency of Cpf1 mutagenesis, the Pearson correlation was calculated between each permutation pair in terms of log_2_ rpm abundance. This value was defined as the permutation correlation. **b.** Empirical cumulative density plot of all permutation correlations across the 4,704 crRNA array combinations in the CCAS library. Greater than half of all crRNA array combinations had a correlation coefficient R ≥ 0.97, indicating that the majority of crRNA array permutations were strongly correlated. **c.** Scatterplot comparing log_2_ rpm abundance of crH2-Q2.1_crPten.240 and its permutation crPten.240_crH2-Q2.1 across all 10 tumor samples. The correlation coefficient and associated p-value of the correlation are noted in the top left (R = 0.999, *p* = 2.28 e-19). **d.** Scatterplot comparing log_2_ rpm abundance of crCbwd1.84_crEpha2.5 and its permutation crEpha2.5_crCbwd1.84 across all 10 tumor samples. The correlation coefficient and associated p-value of the correlation are noted in the top left (R = 0.999, *p* = 7.09 e-19). **e.** Marginal distribution meta-analysis of all 98 constituent single crRNAs in the CCAS library showing the average log_2_ rpm abundance of all DKO crRNA arrays associated with each individual crRNA when present in position 1 or in position 2 of the crRNA array. The scatterplot shows the average log_2_ rpm abundance for each single crRNA when in position 1 (x-axis) or position 2 (y-axis). Across all 98 single crRNAs, the average abundance for each single crRNA when in position 1 was significantly correlated with the average abundance when in position 2 (Pearson correlation coefficient (R) = 0.397, *p* = 5.25 e-5 by t-distribution), showing that individual crRNAs confer a similar selective advantage regardless of position in the crRNA array.

**Supplementary Figure S5:**
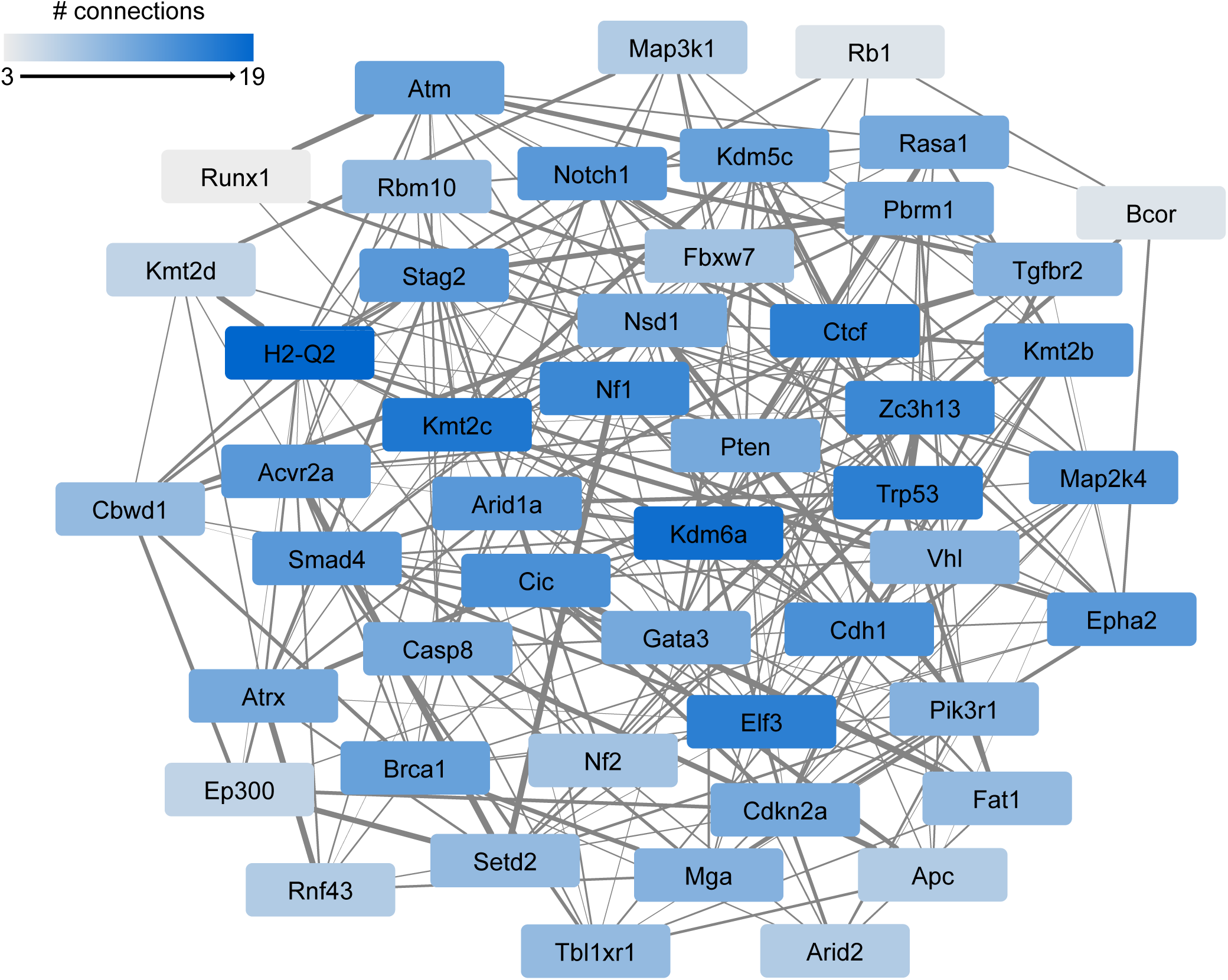
Complete network analysis of synergistic driver pairs. The complete map of the gene-level synergistic driver network among all 49 genes in the CCAS library is shown. Each node represents one gene, and each edge indicates a statistically significant synergistic interaction between a given gene pair (Benjamini-Hochberg adjusted p-value < 0.05, as in **Fig. 4b**). The strength of each synergistic interaction (SynCo score) is represented by edge width. Nodes are color-coded based on the degree of connectivity within the network.

**Supplementary Figure S6:**
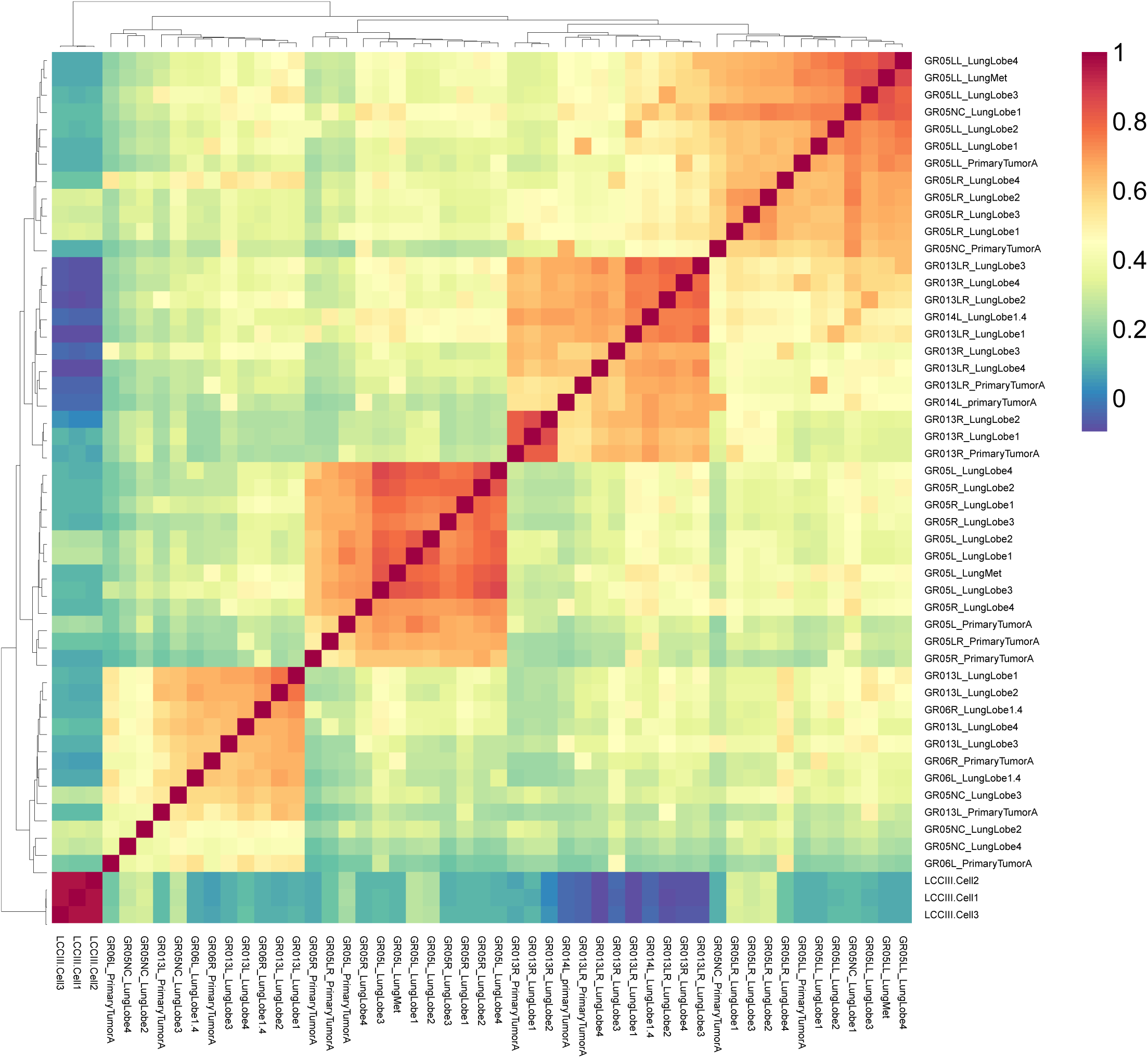
Heatmap of pairwise Pearson correlation coefficients of crRNA arrays in the CCAS metastasis screen. Heatmap of pairwise Pearson correlation coefficients in log_2_ rpm abundance from all 50 samples, including CCAS transduced cells before transplantation (day 7 post infection, n = 3 biological replicates), primary tumors (n = 11 tumors from 11 mice, 7 were *Nu/Nu* and 4 were *Rag1*-/-), and metastases (n = 36 samples from 11 mice).

**Supplementary Figure S7:**
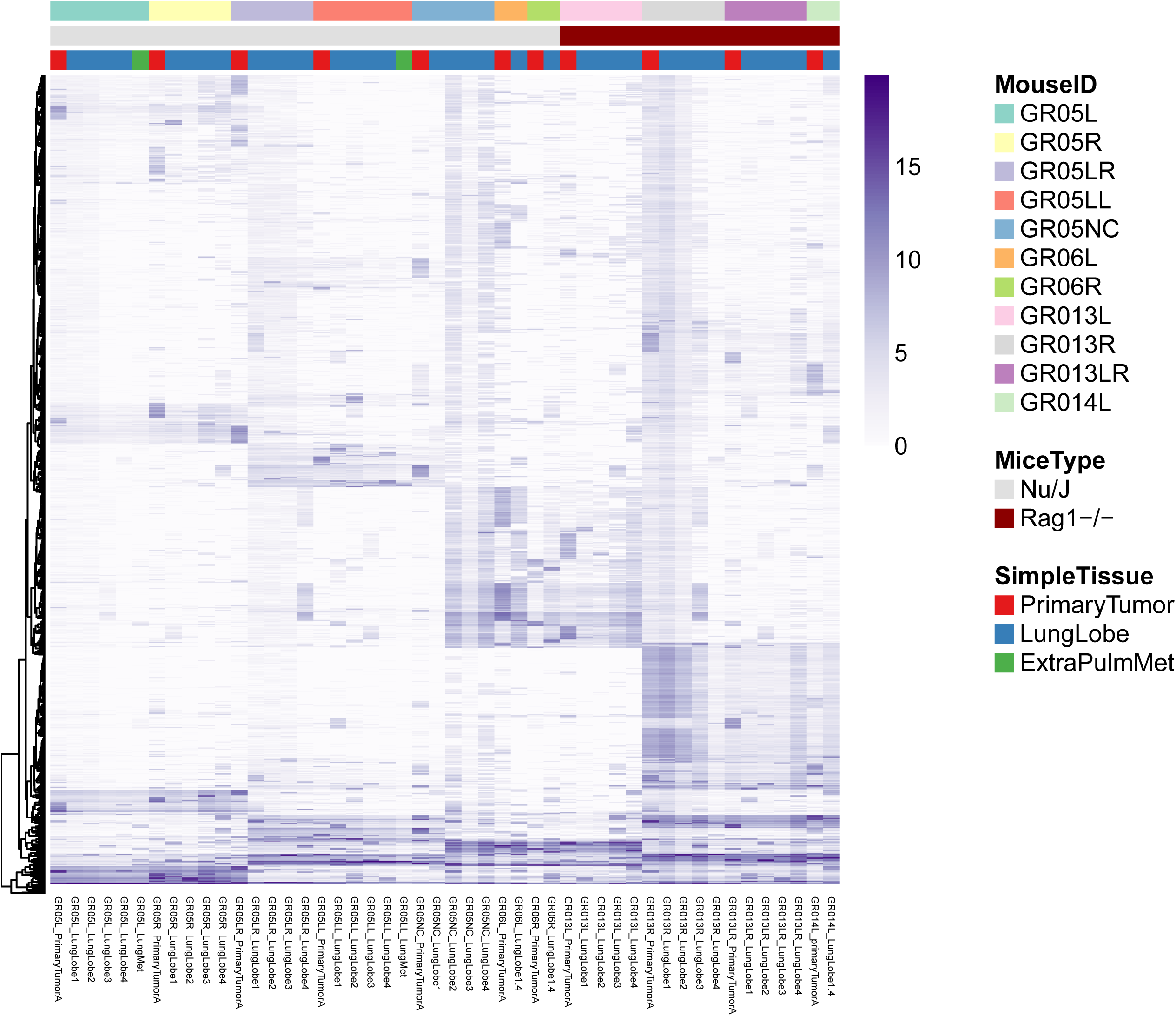
Overall library representation landscape of all crRNA array abundance in the CCAS metastasis screen. Heatmap of all crRNA array abundance in log_2_ rpm abundance from all 50 samples, including CCAS transduced cells before transplantation (day 7 post infection, n = 3 biological replicates), primary tumors (n = 11 tumors from 11 mice, 7 were *Nu/Nu* and 4 were *Rag1*-/-), and metastases (n = 36 samples from 11 mice).

**Supplementary Figure S8.**
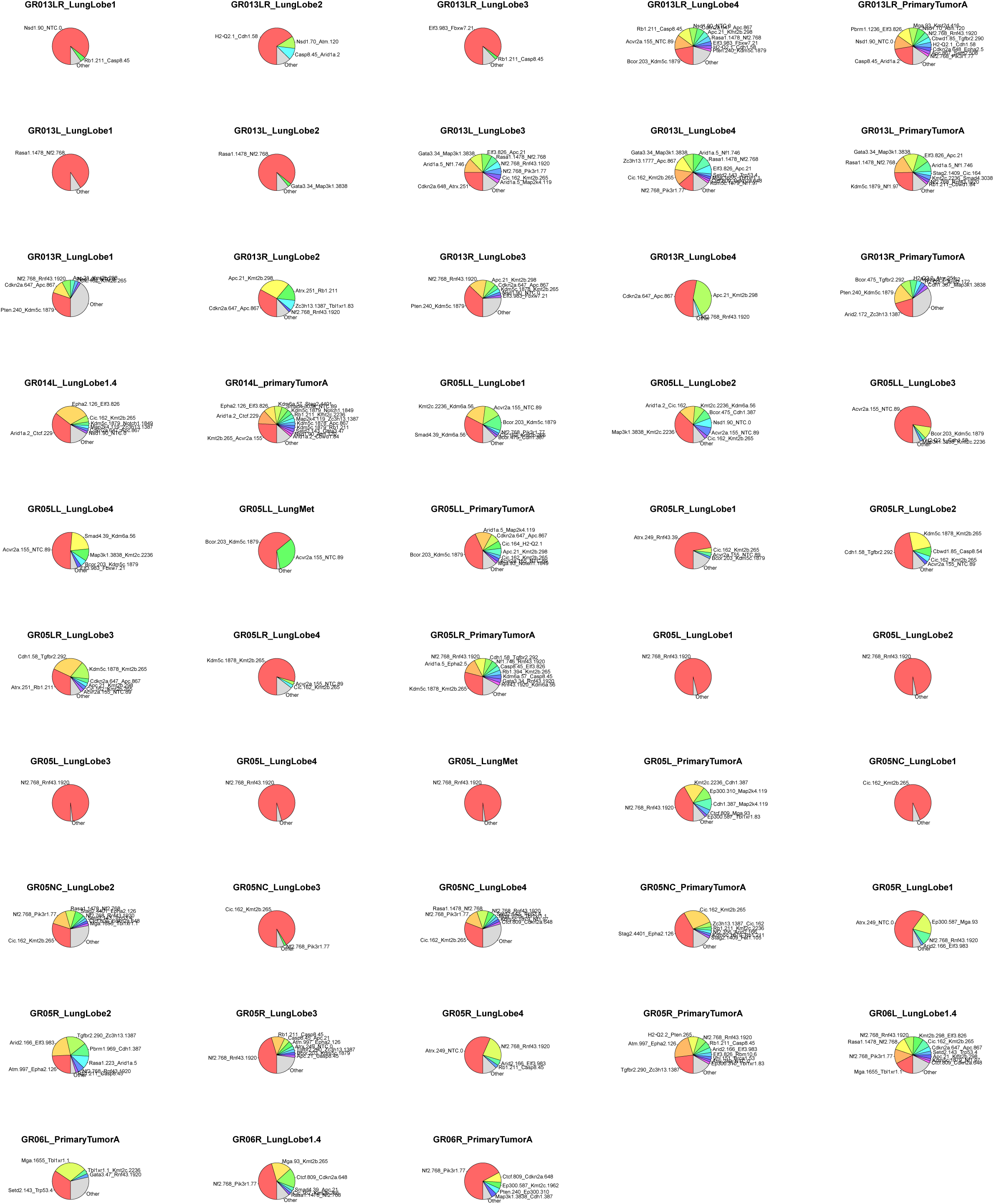
Pie charts of dominant clones in all primary tumor and metastases in the CCAS metastasis screen. Pie charts showing dominant crRNA arrays (>2% reads) in each sample, across all 11 primary tumors and 36 metastasis samples. The area for each crRNA array corresponds to the percentage of reads within the tumor.

## Supplementary Tables

**Table S1:** Ranked list of putative TSGs from analysis of 17 cancer types from TCGA (PANCAN17-TSG50).

**Table S2:** Spacer and oligo sequences of all crRNA arrays represented in the PANCAN17-mTSG CCAS library.

**Table S3:** Raw abundance of all CCAS crRNA arrays in the transformation-tumorigenesis screen.

**Table S4:** Log2 rpm abundance of all CCAS crRNA arrays in the transformation-tumorigenesis screen.

**Table S5:** Number of detected crRNA array counts in plasmid, cells, and tumor samples in the transformation-tumorigenesis screen.

**Table S6:** Significant outlier crRNA arrays in individual tumors compared to cells in the transformation-tumorigenesis screen.

**Table S7:** Significantly enriched crRNA arrays in tumors, compared to NTC-NTC controls in the transformation-tumorigenesis screen.

**Table S8:** Number of significant crRNA arrays associated with each target gene in the transformation-tumorigenesis screen.

**Table S9:** Number of significant DKO crRNA arrays associated with each gene pair in the transformation-tumorigenesis screen.

**Table S10:** Marginal distribution analysis of single crRNAs in position 1 and in position 2 of the crRNA array in the transformation-tumorigenesis screen.

**Table S11:** Permutation correlation analysis of all DKO crRNA array combinations in the CCAS library in the transformation-tumorigenesis screen.

**Table S12:** SynCo scores for each crRNA array in each tumor sample in the transformation-tumorigenesis screen.

**Table S13:** Statistical analysis of SynCo scores for each crRNA array across all tumor samples in the transformation-tumorigenesis screen.

**Table S14:** Co-mutation analysis of all synergistic driver pairs across 21 human cancer types.

**Table S15:** Number of cancer types in which Synco-synergistic pairs in the transformation-tumorigenesis screen that are significantly co-mutated.

**Table S16:** Metadata of the CCAS metastasis screen.

**Table S17:** Raw counts of all crRNA arrays in the metastasis screen.

**Table S18:** Log_2_ normalized counts of all crRNA arrays in the metastasis screen.

**Table S19:** Significantly enriched crRNA arrays compared to NTC-NTC controls of all primary tumors and metastases in the metastasis screen.

**Table S20:** Rank list of genes associated with significantly enriched crRNA arrays compared to NTC-NTC controls of all primary tumors and metastases in the metastasis screen.

**Table S21:** Rank list of gene pairs by number of independent crRNA arrays significantly enriched compared to NTC-NTC controls of all primary tumors and metastases in the metastasis screen.

**Table S22:** Marginal distribution analysis of single crRNAs in position 1 and in position 2 of the crRNA array in the metastasis screen.

**Table S23:** Analysis of metastasis-specific dominant clones of crRNA arrays in the metastasis screen.

## Acknowledgments

### Support

We thank all members in the Chen laboratory, as well as various colleagues in Department of Genetics, Immunobiology Program, Systems Biology Institute, Cancer Center and Stem Cell Center at Yale for assistance and/or discussion. We thank the Center for Genome Analysis, Center for Molecular Discovery, High Performance Computing Center, West Campus Analytical Chemistry Core and West Campus Imaging Core and Keck Biotechnology Resource Laboratory at Yale, for technical support. SC is supported by Damon Runyon Research Foundation Dale Frey Award for Breakthrough Scientists (DRG-2117-12; DFS-13-15), Melanoma Research Foundation (MRA-412806), St-Baldrick’s Foundation (426685), Yale Institutional Research Grant from American Cancer Society (IRG 58-012-54), Breast Cancer Alliance, Yale Skin Cancer SPORE (A08306), Yale Lung Cancer SPORE (A10805) and NIH/NCI Center for Cancer Systems Biology (U54). RDC is supported by the Yale MSTP training grant from NIH (T32GM007205). AC is supported by Yale PhD training grant from NIH (T32GM007223). GW is supported by RJ Anderson Postdoctoral Fellowship.

### Compliance and approval

All animal work was performed under the guidelines of Yale University Institutional Animal Care and Use Committee (IACUC) with approved protocol (Chen-2015-20068), and complied with the Guide for Care and Use of Laboratory Animals, National Research Council, 1996 (institutional animal welfare assurance no. A-3125-01).

### Contributions

RDC designed and generated the vectors, libraries and initiated the study. GW performed majority of cellular, animal experiments and screen readout. AC developed, optimized the screening system, and performed cellular and animal experiments. LY assisted with part of experiments. RDC developed statistical algorithms and computational pipelines, analyzed screen data and prepared figures. RDC and SC wrote the paper with inputs from GW and <u>AC. SC</u> conceived the study, provided conceptual advice and supervised the work.

## Methods

### Animal work statements

All animal work was performed under the guidelines of Institutional Animal Care and Use Committee (IACUC) with an approved protocol, and complied with the Guide for Care and Use of Laboratory Animals, National Research Council, 1996 (institutional animal welfare assurance no. A-3125-01).

### Design, synthesis and cloning of the CCAS library

Significantly mutated genes (SMGs) were identified by analysis of pan-cancer mutation data of 17 cancer types from The Cancer Genome Atlas downloaded via Synapse (https://www.synapse.org/#!Synapse:syn1729383) and from the Broad Institute GDAC (http://gdac.broadinstitute.org/). The top 50 putative TSGs were chosen in an unbiased manner using a multistep approach that prioritizes genes, which are significantly mutated in multiple cancer types and possess mutational signatures consistent with non-oncogenes. **1)** We first compiled a list of all significantly mutated genes in each of the 17 cancer types by collecting all MutSig2CV results from GDAC and using a cutoff of q < 0.1. **2)** To remove putative oncogenes from the significantly mutated gene sets in each cancer type, we calculated the ratio of null to silent mutations for each SMG in that cancer, and multiplied this ratio by the square root of the number of null mutations. **3)** Ratio scores for each gene were then summed across cancer types. **4)** Finally, to heavily weight genes that are SMGs in multiple cancer types, we multiplied the summed ratio scores by the number of unique cancer types in which a gene was considered an SMG. The resulting gene set is defined as PANCAN17-TSG50.

Of the top 50 putative TSGs identified by this approach, 49 were found to have clear mouse orthologs (defined as PANCAN17-mTSG). We then analyzed the complete exon sequences of these 49 genes to extract all possible Cpf1 spacers (i.e., all 20mers beginning with the Cpf1 PAM, 5’-TTTN-3’). Each of these 20mers was then reverse complemented and mapped to the entire mm10 reference genome by Bowtie 1.1.2 (*80*), with settings bowtie -n 2 -l 18 -p 8 -a -y --best -e 90. After filtering out all alignments that contained mismatches in the final 3 basepairs (corresponding to the Cpf1 PAM) and disregarding any mismatches in the fourth to last basepair, we quantified the number of genome-wide alignments for each crRNA using all 0, 1, and 2 mismatch (mm) alignments. A total mismatch score (MM score) was calculated for each crRNA using the following ad hoc formula: MM score = 0mm*1000 + 1mm*50 + 2mm*1. An “on-target” (OT) score was also approximated by counting the number of consecutive thymidines in each crRNA, and then using the formula: OT score = 100 / (max_consecutive_Thymidines)^2^. We then sorted all the crRNAs corresponding to each target gene by low MM score and high OT score. Finally, the top 2 crRNAs for each gene were chosen. In the event of ties, crRNAs targeting constitutive exons and/or the first exon were prioritized. 3 NTC crRNAs were randomly generated.

To generate the 9,408 DKO crRNA arrays in the library, we first computed all possible permutations of the 98 gene-targeting crRNAs, with the stipulation that crRNAs targeting the same gene would not be included in the same crRNA array. For SKO crRNA arrays, we placed each gene-targeting crRNA in the first position of the crRNA array and toggled the 3 NTCs through the second position (98 * 3 = 294 crRNA arrays). Finally, we generated 3 NTC-NTC crRNA arrays from various combinations of the 3 NTC single crRNAs.

### Cell lines

A non-small cell lung cancer (NSCLC) cell line ^21^ (KPD cell line) was used for initial testing of crRNA array constructs. An immortalized, but non-transformed hepatocyte cell line (clone IM) was transduced with LentiCpf1 to generate Cpf1-positive cells (IM.C9-Cpf1). All cell lines were grown under standard conditions using DMEM containing 10% FBS, 1% Pen/strep in a 5% CO2 incubator.

### Nextera analysis of indels generated by Cpf1

CrRNA arrays (crPten.crNf1 and crNf1.crPten) were cloned into Lenti-U6-crRNA vector, and virus was generated for transduction of KPD cell line.

crPten = TGCATACGCTATAGCTGCTT

crNf1 = TAAGCATAATGATGATGCCA

Seven days after transduction and puromycin selection, genomic DNA from harvested from the cells in culture. The surrounding genomic regions flanking the target sites of crPten and crNf1 were first amplified by PCR using the following primers (5’– 3’): Pten_fwd = ACTCACCAGTGTTTAACATGCAGGC, Pten_rev = GGCAAGGTAGGTACGCATTTGCT; Nf1_fwd = AGCAGCTGTCCTGGCTGTTC, Nf1_rev = CGTGCACCTCCCTTGTCAGG. Nextera XT library preparation was then performed according to manufacturer protocol. Reads were mapped to the mm10 mouse genome using BWA ^70^, with the settings bwa mem -t 8 -w 200. Indel variants were first processed with Samtools ^71^ with the settings samtools mpileup -B -q 10 -d 10000000000000, then piped into VarScan v2.3.9 ^72^ with the settings pileup2indel --min-coverage 1 --min-reads2 1 --min-var-freq 0.00001.

### Lentiviral library production

The LentiCpf1, Lenti-U6-crRNA vector and Lenti-CCAS library plasmids were used to make vector or library-containing lentiviruses. Briefly, envelope plasmid pMD2.G, packaging plasmid psPAX2, and LentiCpf1, Lenti-U6-crRNA or Lenti-CCAS-library plasmid were added at ratios of 1:1:2.5, and then polyethyleneimine (PEI) was added and mixed well by vortexing. The solution was stand at room temperature for 10-20 min, and then the mixture was dropwisely added into 80-90% confluent HEK293FT cells and mixed well by gently agitating the plates. Six hours post-transfection, fresh DMEM supplemented with 10% FBS and 1% Pen/Strep was added to replace the transfection media. Virus-containing supernatant was collected at 48 h and 72 h post-transfection, and was centrifuged at 1500 g for 10 min to remove the cell debris; aliquoted and stored at -80°C. Virus was titrated by infecting IM-Cpf1 cells at a number of different concentrations, followed by the addition of 2 μg/mL puromycin at 24 h post-infection to select the transduced cells. The virus titers were determined by calculating the ratios of surviving cells 48 or 72 h post infection and the cell count at infection.

### CCAS in a mouse model of transformation and early tumorigenesis

Cells were transduced at library transduction was performed with four infection replicates at high coverage and low MOI. Briefly, according to the viral titers, CCAS library lentiviruses were added into a total of > 1×10^8^ IM.C9-Cpf1 cells at calculated MOI of <= 0.2 and incubated 24 h before replacing the viruses-contaning media with 3 μg/mL puromycin containing fresh media to select the virus-transduced cells. Approximately 2×10^7^ cells confer a ~2,000x library coverage. Vector and CCAS library-transduced cells were culture under the pressure of 3 μg/mL puromycin for 7 days before injection or cryopreservation.

Vector and CCAS library-transduced IM.C9-Cpf1 cells were injected subcutaneously into the right and left flanks of Nu/Nu mice at 4×10^6^ cells per flank (~400x coverage per transplant). Tumors were measured every week by caliper and their sizes were estimated as spheres. Statistical significance was assessed by paired t-test.

### Mouse tumor dissection and histology

Mice were sacrificed by carbon dioxide asphyxiation followed by cervical dislocation. Tumors and other organs were manually dissected, and then fixed in 10% formalin for 24-96 hours, and transferred into 70% Ethanol for long-term storage. The tissues were embedded in paraffin, sectioned at 5 μm and stained with hematoxylin and eosin (H&E) for pathological analysis. For tumor size quantification, H&E slides were scanned using an Aperio digital slidescanner (Leica). For molecular biological analysis, tissues were flash frozen with liquid nitrogen, and ground in 5 mL Frosted polyethylene vial set (2240-PEF) in a 2010 GenoGrinder machine (SPEXSamplePrep). Homogenized tissues were used for DNA/RNA/protein extractions.

### CCAS in a mouse model of metastasis

For Cpf1 crRNA array library screen in a mouse model of metastasis, lentiviral pools were generated from the CCAS plasmid library, and transduced ≥ 1×10^8^ Cpf1+ KPD cells with three independent infection replicates at calculated MOI of ≤ 0.2 and incubated 24 h before replacing the viruses-contaning media with 3 μg/mL puromycin containing fresh media to select the virus-transduced cells. Approximately 2×10^7^ cells confer a ~2,000x library coverage. CCAS library-transduced cells were culture under the pressure of 3 μg/mL puromycin for 7 days before injection or cryopreservation.

CCAS-treated cells were then injected at 4×10^6^ cells per mouse (~400x coverage) subcutaneously into *Nu/Nu* mice (n = 7) and *Rag1*-/- mice (n = 4). Metastases were allowed to form *in vivo* for 8 weeks after injection. Primary tumors, four lung lobes, and other stereoscope-visible metastases, were then dissected and then subjected to genomic DNA extraction and crRNA array sequencing.

### Genomic DNA extraction

200-800 mg of frozen ground tissue were re-suspended in 6 mL of NK Lysis Buffer (50 mM Tris, 50 mM EDTA, 1% SDS, pH 8.0) supplemented with 30 μL of 20 mg/mL Proteinase K (Qiagen) in 15 mL conical tubes, and incubated at 55°C bath for 2 h up to overnight. After all the tissues have been lysed, 30 μL of 10 mg/mL RNAse A (Qiagen) was added, mixed well and incubated at 37°C for 30 min. Samples were chilled on ice and then 2 mL of pre-chilled 7.5 M ammonium acetate (Sigma) was added to precipitate proteins. The samples were inverted and vortexed for 15-30 s and then centrifuged at ≥ 4,000 *g* for 10 min. The supernatant was carefully decanted into a new 15 mL conical tube, followed by the addition of 6 mL 100% isopropanol (at a ratio of ~ 0.7), inverted 30-50 times and centrifuged at ≥ 4,000 *g* for 10 minutes. Genomic DNA should be visible as a small white pellet. After discarding the supernatant, 6 mL of freshly prepared 70% ethanol was added, mixed well, and then centrifuged at ≥ 4,000 *g* for 10 min. The supernatant was discarded by pouring; and remaining residues was removed using a pipette. After air-drying for 10-30 min, DNA was re-suspended by adding 200-500 μL of Nuclease-Free H_2_O. The genomic DNA concentration was measured using a Nanodrop (Thermo Scientific), and normalized to 1000 ng/μL for the following readout PCR.

### Cpf1 CrRNA array library readout

The crRNA array library readout was performed using a 2-step PCR approach. Briefly, in the 1st round PCR, enough genomic DNA was used as template to guarantee coverage of the library abundance and representation. For example, assuming 6.6 pg of gDNA per cell, 20-48 μg of gDNA (≥75x) was used per sample. For the 1st PCR, the sgRNA-included region was amplified using primers specific to the double-knockout CCAS vector using Phusion Flash High Fidelity Master Mix (ThermoFisher) with thermocycling parameters: 98°C for 1 min, 15 cycles of (98°C for 1s, 60°C for 5s, 72 °C for 15s), and 72 °C for 1 min. Fwd AATGGACTATCATATGCTTACCGTAACTTGAAAGTATTTCG Rev CTTTAGTTTGTATGTCTGTTGCTATTATGTCTACTATTCTTTCCC

In the 2nd PCR, 1^st^ round PCR products for each biological repeats were pooled, then 1-2 μL well-mixed 1st PCR products were used as the template for amplification using sample-tracking barcode primers with thermocycling conditions as 98°C for 1 min, 15 cycles of (98°C for 1s, 60°C for 5s, 72 °C for 15s), and 72 °C for 1 min. The 2^nd^ PCR products were quantified in 2% E-gel EX (Life Technologies) using E-Gel^®^ Low Range Quantitative DNA Ladder (ThermoFisher), then the same amount of each barcoded samples were combined. The pooled PCR products were purified using QIAquick PCR Purification Kit and further QIAquick Gel Extraction Kit from 2% E-gel EX. The purified pooled library was quantified in a gel-based method. Diluted libraries with 5-20% PhiX were sequenced with Hiseq 2500 or HiSeq 4000 systems (Illumina) with 150bp paired-end read length.

### Cpf1 double knockout Illumina data pre-processing

Raw single-end fastq read files were filtered and demultiplexed using Cutadapt ^73^. To remove extra sequences downstream (i.e. 3’ end) of the dual-RNA spacer sequences, including the U6 terminator, we used the following settings: cutadapt --discard-untrimmed –e 0.1 -a TTTTTTAAGCTTGGCGTGGATCCGATATCA. As the forward PCR primers used to readout crRNA array representation were designed to have a variety of barcodes to facilitate multiplexed sequencing, we then demultiplexed these filtered reads with the following settings: cutadapt -g file:fbc.fasta --no-trim, where fbc.fasta contained the 12 possible barcode sequences within the forward primers. Finally, to remove extraneous sequences upstream (i.e. 5’ end) of the crRNA array spacers, including the first DR, we used the following settings: cutadapt --discard-untrimmed –e 0.1 –g AAAGGACGAAACACCgTAATTTCTACTAAGTGTAGAT. Through this procedure, the raw fastq read files were pared down to the sequences of the first crRNA, the second DR, and finally the second crRNA (cr1-DR-cr2). The filtered fastq reads were then mapped to the CCAS reference index. To do so, we first generated a bowtie index of the CCAS library using the bowtie-build command in Bowtie 1.1.2 ^69^. Using these bowtie indexes, we mapped the filtered fastq read files using the following settings: bowtie -v 2 -k 1 -m 1 --best. These settings ensured only single-match reads would be retained for downstream analysis.

### Analysis of CCAS library representation

Using the resultant mapping output, we quantitated the number of reads that had mapped to each crRNA array within the library. We normalized the number of reads in each sample by converting raw crRNA array counts to reads per million (rpm). The rpm values were then subject to log_2_ transformation for certain analyses. To generate correlation heatmaps, we used the *NMF* R package. To generate sgRNA representation barplots, we set a detection threshold of log_2_ rpm ≥ 1, and counted the number of unique crRNA arrays present in each sample.

### Analysis of enriched DKO and SKO crRNA arrays

To directly compare the abundance in tumor samples vs. cells, we performed linear regression and identified significant outliers using the *outlierTest* function from the *car* R package. Significant outlier crRNA arrays in individual tumors vs. cells were defined as having a Bonferroni adjusted *p* < 0.05, based on analysis of the studentized regression residuals.

To identify crRNA arrays significantly enriched above NTC-NTC controls, we similarly performed two-sided t-tests on the log2 rpm abundance of each crRNA array compared to the average of all NTC-NTC crRNA arrays. Significantly enriched crRNA arrays were defined as having a Benjamini-Hochberg adjusted *p* < 0.05 Each significantly enriched crRNA array was then deconstructed into its two constituent crRNAs, and finally down to the two target genes. We used this 3-tiered dataset to determine how many genes were involved in an enriched crRNA array (either SKO or DKO). Finally, we compiled all of the significant crRNA arrays associated with each gene, and counted the number of DKO or SKO crRNA arrays.

### Position effect analysis of crRNA permutations

Marginal distribution analysis was performed by considering each of the 98 single crRNAs when found in position 1 or position 2 of the crRNA array. Specifically, the average log_2_ rpm abundance was calculated for each single crRNA, and these average scores were compared between position 1 and position 2. For direct permutation correlation analysis, we condensed the 9,408 DKO crRNA arrays down into 4,704 crRNA array combinations (i.e., crX.crY and crY.crX are two permutations of the same combination). We then calculated the correlation between the two corresponding permutations across all 10 tumor samples (defined as permutation correlation), and assessed the statistical significance by t-distribution. Violin plots, empirical density plots, and scatterplots were generated using these permutation correlation coefficients.

### Synergy analysis of gene pairs

We defined the synergy coefficient (SynCo) for each DKO crRNA array with the following formula:

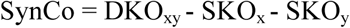

The DKO_xy_ score is the log_2_ rpm abundance of the DKO crRNA array (i.e., crX.crY) after subtracting average NTC-NTC abundance, while SKO_x_ and SKO_y_ scores are defined as the average log_2_ rpm abundance of each SKO crRNA array (3 SKO crRNA arrays associated with each individual crRNA), each after subtracting average NTC-NTC abundance. By this definition, a SynCo score >> 0 would indicate that a given DKO crRNA array is synergistic, as the DKO score would thus be greater than the sum of the individual SKO scores. We calculated the SynCo of each DKO crRNA array within each tumor sample and assessed whether the SynCo score of a given crRNA array across all 10 tumors was statistically significantly different from 0 by a two-sided one-sample t-test. We set a significance threshold of Benjamini-Hochberg adjusted *p* < 0.05, and all significant DKO crRNA arrays with an average SynCo > 0 were considered to be synergistic.

### Network analysis

Using the synergistic crRNA arrays identified through SynCo analysis, we then constructed library-wide networks using individual genes as nodes and SynCo scores as edge weights. The pairwise connections were visualized through Cytoscape 3.4.0 ^74^. Edge width was scaled according to SynCo score. For the global network, node color was additionally scaled according to the degree of network connectivity.

### Analysis of co-mutation patterns in human pan-cancer datasets

For the synergistic driver pairs identified by the CCAS screen, we performed co-mutation analysis on 21 different solid tumor types, all of which were from TCGA except for small cell lung cancer ^75–92^. We obtained the somatic mutation and copy number status of each cohort from cBioPortal ^93,94^ (only somatic mutations were available for lung small cell cancer) and classified all tumors as a mutant or non-mutant for the genes represented in the CCAS library. We defined “mutant” as the presence of nonsynonymous mutations and/or deep deletions in a given gene. After classifying every patient in terms of mutant status, we performed co-mutation (co-occurrence) analysis by calculating the co-occurrence rate for each gene pair. The co-occurrence rate is defined as the intersection (the number of double mutant samples) divided by the union (the number of all single and double mutant samples). Statistical significance was tested by a hypergeometric test, with a significance threshold of Benjamini-Hochberg adjusted *p* < 0.05.

### Analysis of metastasis enrichment over primary tumor and metastatic clonal spread

Comparison of the crRNA array representations was made between metastases to primary tumors. A crRNA array was called metastasis-enriched if it was a dominant clone in a lung lobe or extra-pulmonary metastasis (≥ 2% total reads) but not a dominant clone in the corresponding primary tumor of the same mouse. Waterfall plot was made for all crRNA arrays enriched in a metastases vs primary tumor, ranked by numbers of mice where an crRNA was called enriched.

Monoclonal spread was defined where dominant metastases in all lobes were derived from identical crRNA arrays, and polyclonal spread was defined where dominant metastases in all lobes were derived from multiple varying crRNAs.

#### Blinding statement

Investigators were blinded for sequencing data analysis, but not blinded for tumor engraftment, organ dissection and histology analysis.

#### Code availability

Key scripts used to process and analyze the data will be available to academic community.

#### Accession

Genomic sequencing data will be deposited to NCBI SRA.

#### Key resource sharing

Normalized screen data, sequences of oligos are described in the methods section or supplementary material of the manuscript. Key plasmids and libraries will be deposited to Addgene.

